# Innate sensorimotor policies constrain cerebellar adaptive correction

**DOI:** 10.64898/2026.10.01.754451

**Authors:** Amey Shenoy, Ruben Portugues

## Abstract

Innate sensorimotor policies enable rapid and reliable behavior but can become ineffective when environments change. How these policies interact with feedback-driven processes to support behavioral flexibility remains un-clear. Using closed-loop virtual reality in larval zebrafish, we reversed the visual consequences of self-generated turns. Larvae initially followed their innate mapping and then rapidly switched to corrective turns in the opposite direction. Purkinje cell ablation abolished this correction while preserving innate optomotor responses. Imaging revealed outcome-invariant sensorimotor representations in granule cells and an outcome-dependent modulation of inferior olive neurons and Purkinje cells. Correction weakened as the optic-flow angle increased, despite persistent error-related representations. The innate behavioral repertoire also shifted as the optic-flow angle increased, making corrective turns less likely. An actor-critic model initialized with the measured behavioral repertoire reproduced the limits of correction. Together, these results show that adaptive behavior emerges from the interaction between an innate feed-forward policy and an olivo-cerebellar instructive signal, revealing innate behavioral structure as a fundamental constraint on cerebellar adaptation.

## Introduction

Animals often encounter situations that require rapid behavioral responses. Rather than generating actions de novo, they rely on innate sensorimotor policies that transform sensory inputs into reliable motor outputs through evolutionarily conserved circuits [1]. These circuits provide robust behavior but cannot account for every perturbation encountered in natural environments. Thus, animals require mechanisms to continuously evaluate the sensory consequences of their motor actions and use this information to modify future behavior [2].

At an algorithmic level, adaptive behavior has traditionally been studied through supervised and reinforcement learning frameworks, often associated with the cerebellum and basal ganglia, respectively [3]. Cerebellar learning is thought to rely on instructive signals that report deviations between expected and actual sensory consequences [4–7], whereas reinforcement learning uses action outcomes to modify future action selection [8]. However, natural behavior is unlikely to follow such a clear division, as sensory consequences of an action can simultaneously provide both an instructive error and information about whether that action should be repeated. Moreover, growing evidence for reward- and outcome-related signals within cerebellar circuits challenges a strict separation of these computations across brain regions [9–11]. How error-driven and reinforcement-like mechanisms interact to support adaptive behavior in natural settings remains unclear.

The larval zebrafish optomotor response (OMR) provides a powerful system for addressing these questions. During OMR, larval zebrafish turn and swim in the direction of the whole field optic flow in order to stabilize their position in space [12–14]. Although this behavior is largely mediated by feed-forward sensorimotor transformations [15–17], larval zebrafish can modify their behavior in response to altered visual feedback [18–22], opening avenues for examining the circuits underlying action generation and adaptation.

Here, we developed a closed-loop virtual reality paradigm in which the visual consequences of self-generated turns [23] could be precisely controlled. By reversing the sensory feedback [24, 25], we created a visuomotor mismatch in which the innate response via feedforward circuits increased the visual error rather than reducing it. This mismatch serves both as an instructive error and as information for outcome-dependent action selection, linking computations traditionally studied within supervised- and reinforcement-learning frameworks.

We show that larvae rapidly performed corrective turns in the opposite direction to counteract the effects of this visuomotor mismatch while preserving their initial response, demonstrating that adaptive behavior is achieved by superimposing feedback-driven corrections onto an existing feed-forward action strategy rather than by relearning this mapping. Chemogenetic ablations [26] of Purkinje cells revealed a selective requirement for the cerebellar circuit in this correction, while population imaging identified representations of sensory and motor context in granule cells, and action outcomes across inferior olive and Purkinje cell populations. This behavioral correction progressively weakened as the optic flow angle increased despite persistent error-related representations. Instead, the innate behavioral repertoire shifted in an angle-dependent manner, reducing the availability of corrective turns as the optic flow angle increased. Finally, using an actor–critic model [8], we show that this angle-dependent shift in the innate behavioral repertoire is sufficient to reproduce the limits of correction. Together, these findings show that adaptive behavior emerges from the interaction between an innate feed-forward policy and feedback about the consequences of action, and the structure of the existing behavioral policy is a fundamental constraint on error-driven adaptation [27, 28].

## Results

To understand how altered sensory consequences modulate behavior [29, 30], we developed a closed-loop virtual reality paradigm in which head-restrained larval zebrafish were presented with whole-field optic flow moving at a defined angle relative to their body axis (Figure 1a, see Methods). Tail movements were tracked with a highspeed camera and used to update the visual stimulus in real time [31], providing precise control over the visual feedback associated with each turn. We refer to the relationship between a turn and its visual consequence as the visuomotor rule. Under the normal rule, turning in the direction of the optic flow would rotate it towards the fish, progressively reducing the angular offset between the optic flow and the fish. This mimics the feedback larvae would experience while swimming in a natural environment. Under the inverse rule, the relationship between turning and visual feedback is reversed such that turning in the direction of the optic flow would rotate it away from the fish, creating a visuomotor mismatch between the action and the sensory outcome (Figure 1b, Figure S1a). However, it is important to note that the numerical value of the gain was low, resulting in a small angle change in either condition (Figure S1c).

**Figure 1:**
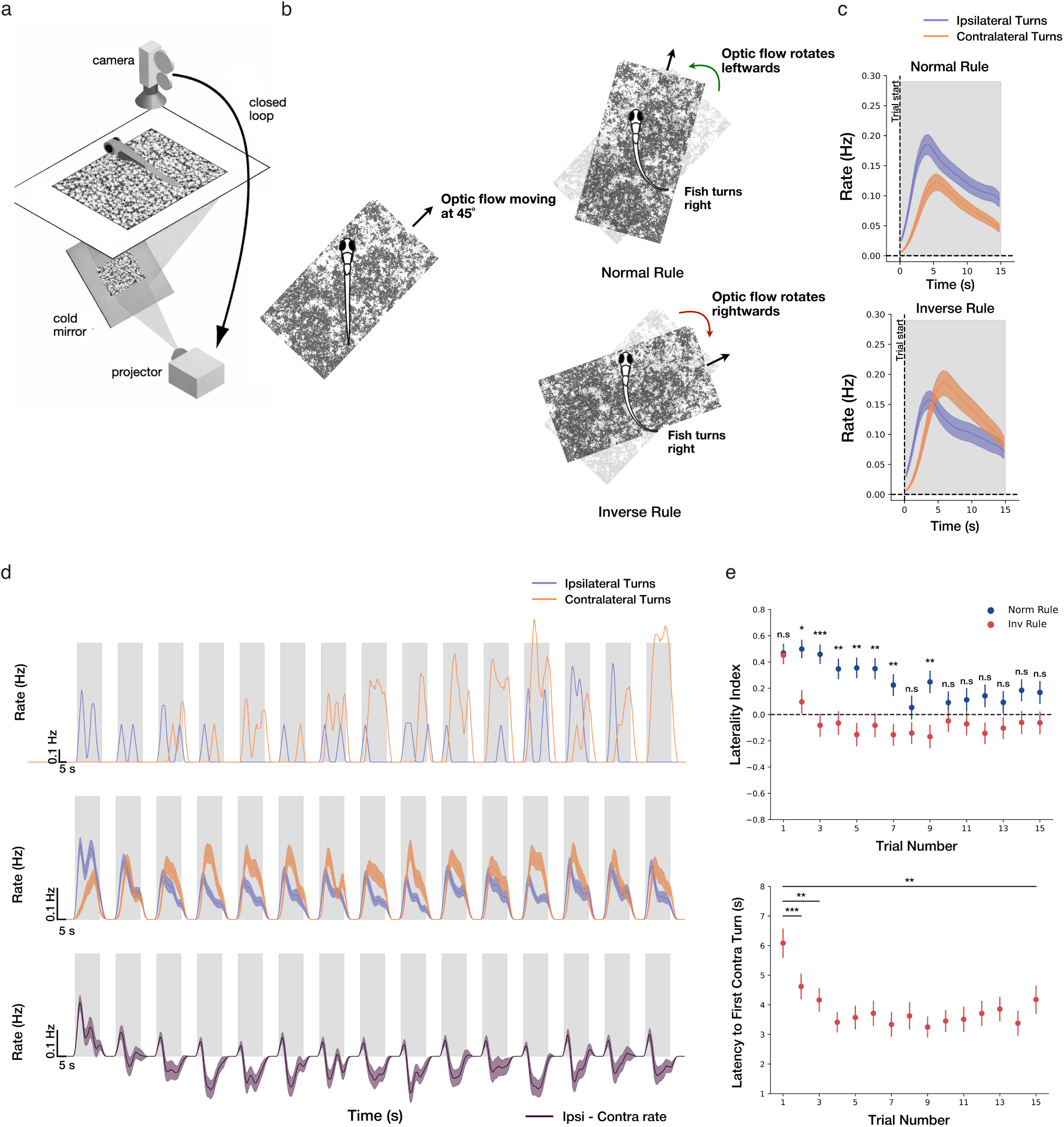
Visuomotor mismatch drives rapid behavioral correction. **a.** Closed-loop virtual-reality setup for head-restrained larvae. Tail movements were tracked and used to update whole-field optic flow in real time. **b.** Visuomotor rules. Optic flow moved rightward (+45°) or leftward (−45°) relative to the body axis. Left-trial turns are sign-inverted so that, throughout, turns are classified as ipsilateral (in the direction of optic flow) or contralateral (opposite to optic flow). Under normal rule, ipsilateral turns rotate the flow toward the body axis (reducing the offset); under inverse rule, the same turns rotate it away (increasing the offset). **c.** Mean ipsilateral and contralateral turn rate, averaged across trials, under normal (top) and inverse (bottom) rule trials. **d.** Trial-wise ipsilateral (blue) and contralateral (red) turn rate for a representative larva (top) and averaged across larvae (middle), and the turning bias Δrate = ipsilateral − contralateral rate averaged across larvae (bottom) for inverse rule trials. **e.** Trial-wise laterality index (LI) under normal and inverse rule trials (top), and latency to the first contralateral turn across inverse-rule trials (bottom). All plots show mean and sem (shaded areas and error bars) across fish (n = 53 fish).

### Visuomotor mismatch drives rapid behavioral correction

When presented with optic flow at 45° relative to the body axis under normal rule, larvae (n = 53) turned predominantly in the direction of the optic flow (ipsilateral turns) (Figure 1c top), rotating it towards the body axis (Figure S1c left), while also producing a substantial fraction of turns in the opposite direction (contralateral turns). Ipsilateral turn rate exceeded contralateral rate (0.12 ± 0.01 versus 0.07 ± 0.008 turns/s Wilcoxon signed-rank test, p < 0.001), yielding a positive turning bias (Δrate = ipsiRate − contraRate = 0.043 ± 0.015 turns/s; one-sample Wilcoxon signed-rank test versus zero p<0.01) (Figure S1b). During inverse rule trials, larvae initially turn in the ipsilateral direction, followed by a rapid increase in the contralateral turning rate (Figure 1c bottom), resulting in a negative Δrate significantly lower compared to the normal rule trials (Δrate = -0.04 ± 0.02 turns/s Wilcoxon signed-rank test p < 0.001), demonstrating a switch to contralateral turns that would bring the optic flow closer to the fish (Figure S1c right). This behavioral shift emerged progressively across trials. In the initial inverse rule trials, larvae followed their innate stimulus-action mapping and turned ipsilaterally (Figure 1d top). However, under inverse rule, these turns rotate the optic flow away from the fish, increasing the visual error (Figure S1c, right). After experiencing this feedback for 2-3 trials, larvae start turning contralaterally, rotating the optic flow back towards them (Figure S1c right).

To quantify this trial-wise behavioral shift, we compute a laterality index(LI) (see Methods) for each trial, which is defined as the normalized difference between the number of ipsilateral and contralateral turns that a fish performs during that trial. Under normal rule, LI remained positive across all trials (Figure 1e top) (meanLI = 0.25 ± 0.05, one-sample Wilcoxon signed-rank test versus 0, p < 0.0001), reflecting an ipsilateral turning bias consistent with the innate stimulus-action mapping. Under inverse rule, LI was initially positive but shifted to negative values within 2-3 trials, demonstrating a rapid transition towards corrective turning behavior (Figure 1e top).

Importantly, this behavior change does not arise through an inversion of the innate stimulus-action mapping. At the start of each trial, larvae consistently follow their innate stimulus-action mapping and turn in the naive ipsilateral direction and then switch to corrective contralateral turns (Figure 1d bottom). To examine whether this initial response changed over the course of adaptation, we compared the distributions of first-turn direction and angle across the first, middle, and last five trials. First-turn distributions remained highly similar across these epochs for both normal- and inverse-rule trials (Figure S1d), indicating that the initial stimulus-evoked response was maintained even as the distribution of subsequent actions changed substantially. Thus, rapid behavioral correction can emerge without an immediate remapping of the initial sensorimotor response. Consistent with this, the latency to the first turn in the contralateral direction decreases from 6.64 ± 0.5 s (trial1) to 4.37 ± 0.4 s (trial15) (Figure 1e bottom), indicating that corrective actions were recruited progressively earlier with experience.

Turn kinematics were also modulated in response to visuomotor rules and sensory outcomes [21]. Larvae show higher vigor turns during normal rule (normVig = 1.16 ± 0.03 vs invVig = 1.06 ± 0.03, Wilcoxon signed-rank test, p < 0.001), but turn duration and time to initiate the next turn remain similar across conditions (Figure S1e) (p= 0.35 and p = 0.39). However, if we categorize the turns by sensory outcomes (Figure S1f, left), we observe that turns resulting in an unfavorable outcome have significantly longer turning duration (unfavDur = 0.49 ± 0.02 vs favDur = 0.42 ± 0.02, Wilcoxon signed-rank test, p < 0.01), and they also take significantly longer to initiate the next turn after experiencing an unfavorable outcome (unfavT = 2.63 ± 0.15 s vs favT = 2.44 ± 0.18 s, Wilcoxon signed-rank test, p < 0.05), as compared to turns with a favorable outcome (Figure S1g).

Together, these results suggest that larvae generate actions using an innate stimulus-action mapping in a feed-forward manner, which are then evaluated through their sensory consequences, thereby generating error signals that can drive rapid outcome-driven modulation of action selection without requiring an immediate inversion of the initial stimulus–action mapping.

### Optic flow angle sets a limit on behavioral correction

The behavioral correction described so far was measured for optic flow moving at 45° relative to the fish. To understand whether this correction depends on stimulus angle, we varied the initial angle of the optic flow (25°, 45°, 60°, and 90°) and measured turning behavior under normal and inverse rules (Figure 2a).

**Figure 2:**
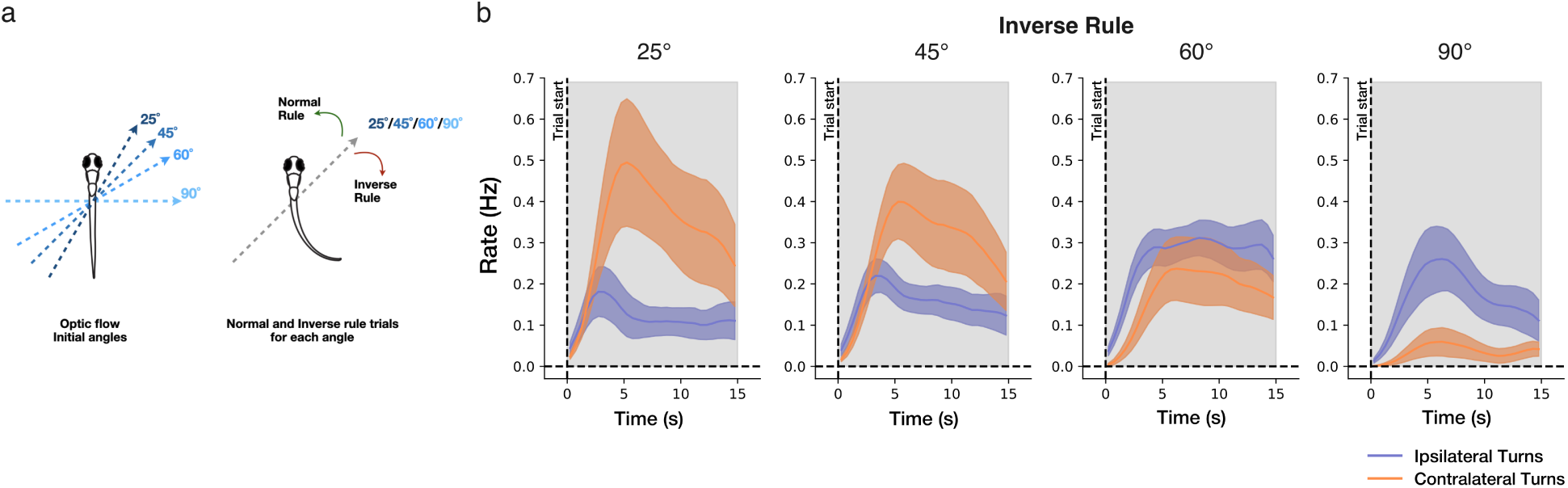
Behavioral correction scales inversely with the angle of the optic flow. **a.** Schematic showing extension of the closed-loop paradigm to four optic flow angles (25°, 45°, 60°, 90°) relative to the body axis. **b.** Mean ipsilateral (blue) and contralateral (orange) turn rate over the 15 s trial, averaged across trials, under inverse rule, for each angle. All plots show mean and sem (shaded areas) across fish (n = 18 fish).

We observed that the behavioral correction systematically weakened as the initial angle increased. Within each 15 s trial, larvae first turned ipsilaterally and then increased the rate of contralateral turns, as shown previously, but the magnitude of the corrective shift depended strongly on the angle (Figure 2b). At 25°, the contralateral rate (0.40 ± 0.12 turn/s) exceeded the ipsilateral turn rate (0.12±0.04 turns/s, p<0.05) within the trial and this rise in contralateral rate become progressively smaller for 45° (0.33 ± 0.08 turns/s), 60° (0.19 ± 0.06 turns/s) and 90° (0.04 ± 0.02 turns/s) where the fish mostly turns ipsilaterally with very few corrective turns (Figure S2b).

Together, these results show that correction is not all-or-none but graded by the stimulus angle. Larvae show strong correction when the optic flow is near the body and progressively less as the angle increases. Having established this behavioral correction and its stimulus dependence, we first asked which circuit produces this correction.

### Purkinje cells are required for correction but not for innate action generation

Larvae do not rely solely on innate stimulus-action mappings to generate rapid and reliable behavior but instead evaluate the sensory consequences of their motor actions to implement corrective actions in response to environmental perturbations (Figure 1). The cerebellum is a candidate circuit for implementing such error-based adaptation [4, 5, 7, 32, 33]. The cerebellar microcircuitry is highly conserved in larval zebrafish [34, 35], with Purkinje cells receiving inputs from granule cells and the inferior olive neurons and relaying them through eurydendroid cells [36] (homologous to the mammalian deep cerebellar nuclei) to downstream sensorimotor regions. This places Purkinje cells at the point where sensory and motor signals are integrated and translated into changes in motor output [37–39].

To test whether cerebellar computations are required for the behavioral corrections we observed in our paradigm, we performed targeted chemogenetic ablation of Purkinje cells (see Methods) and assessed its impact on visuo-motor adaptation under both normal and inverse rule trials.

During normal rule trials, treatment control (unablated, n = 21 fish) and PC-ablated larvae (n=27 fish) showed similar behavior (Figure 3a middle, Figure 3b) with ipsilateral turning rates exceeding contralateral rates in all fish (control ipsi = 0.17 ± 0.02 turn/s vs contra 0.1 ± 0.01 turn/s, Wilcoxon signed-rank test p<0.05 and ablated ipsi = 0.16 ± 0.02 turn/s vs contra 0.1 ± 0.01 turns/s, Wilcoxon signed-rank test p < 0.05). The Δrate remained positive and did not differ between control and ablated groups (0.063 ± 0.02 turns/s vs 0.066±0.02 turns/s Mann-Whitney U test, p = 0.6). The overall turn rate and turn amplitude were similar across groups (Figure S3f, g), indicating that neither the ablation procedure nor the genetic background disrupted the innate optomotor behavior.

**Figure 3:**
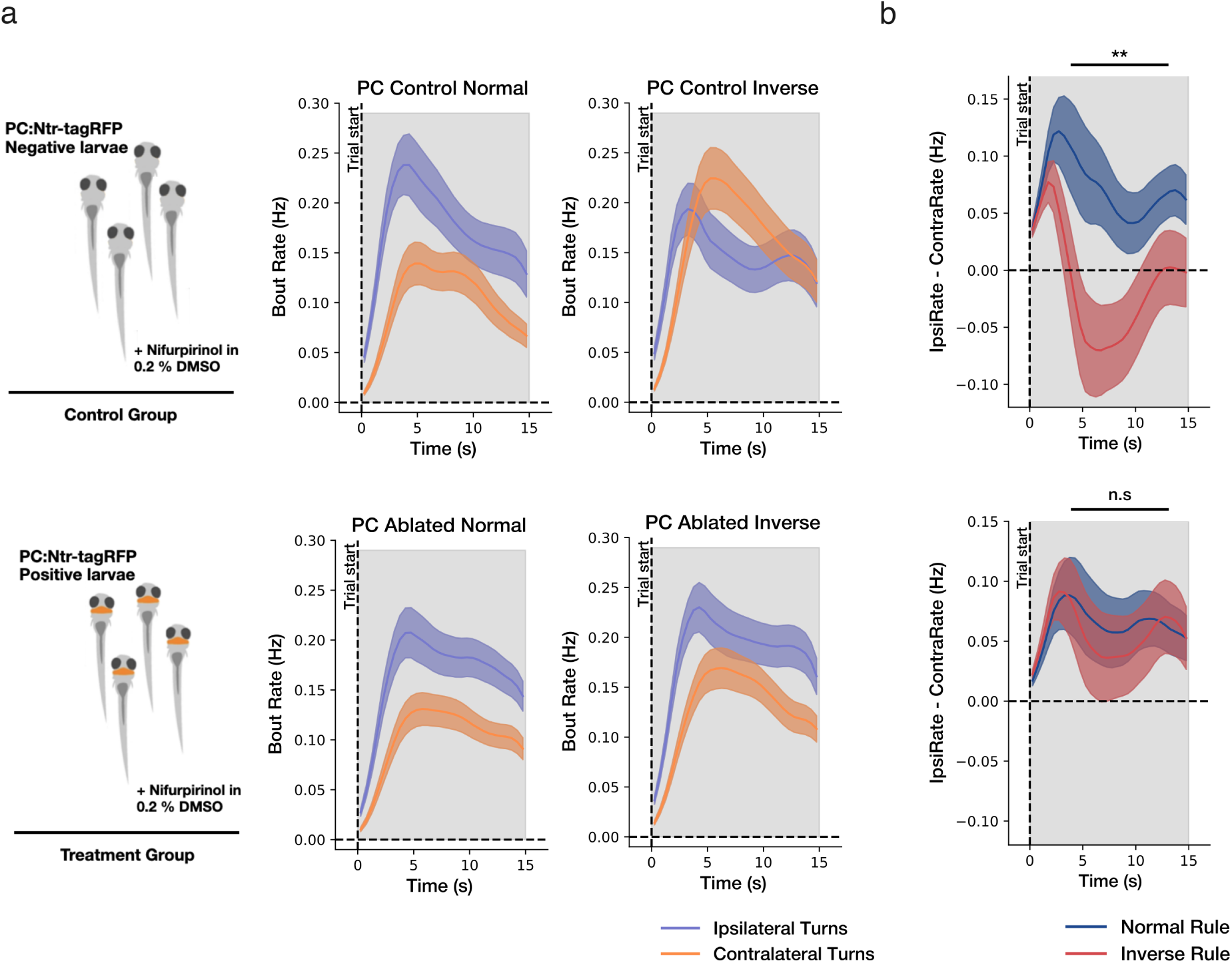
Purkinje cells are required for behavioral correction but not for innate turning. **a.** Top left, control group ([PC:Ntr-tagRFP–negative siblings, identical drug treatment]); bottom left, ablation group ([PC:Ntr-tagRFP–positive larvae, chemogenetically ablated]. Mean ipsilateral and contralateral turn rate, averaged across trials, under normal (middle) and inverse (right) rule for control fish(top) and ablated fish(bottom). **b.** Turning bias Δrate = ipsilateral − contralateral turn rate, averaged across trials, under normal and inverse rule, for control (top) and PC-ablated (bottom) larvae. Horizontal bar above each plot indicates the p-value of the normal-versus-inverse comparison in the 3-12sec time bin (Wilcoxon signed-rank test). All plots show mean and sem (shaded areas) across fish (n = 21 control and 27 ablated fish).

Under inverse rule, the two groups diverged (Figure 3a right, Figure 3b). Control larvae first turned in the ipsilateral direction, and quickly switched to contralateral turns, similar to wildtype fish, resulting in a negative Δrate (normΔrate = 0.07 ± 0.02 turn/s vs invΔrate = -0.04 ± 0.03 turn/s, Wilcoxon signed-rank test p < 0.01). However, PC-ablated larvae did not show this increase in contralateral turning rates as the trial progressed, and the Δrate remained positive for inverse rule trials, similar to normal rule trials (0.06 ± 0.02 turn/s vs 0.05 ± 0.02 turns/s Wilcoxon signed-rank test p = 0.67). Ablated larvae nevertheless showed similar turning rates and turn amplitudes as controls under inverse rule (Figure S3f,g), suggesting that the failure to correct does not arise from a general loss of motor output.

These results show that Purkinje cells are required for feedback-dependent corrections, while the innate optomotor response under normal rule remains largely intact. This suggests that the core feed-forward transformation from optic flow to turning can be expressed without Purkinje-cell output, whereas cerebellar activity becomes particularly important when behavior must be modified according to the sensory consequences of preceding actions.

### Granule cells provide an outcome-invariant sensorimotor basis

Having established that Purkinje cells are necessary for feedback-dependent behavioral adaptations (Figure 3), we asked how the signals that support these modifications are represented in the cerebellar circuitry. We therefore started by performing 2-photon calcium imaging in the main cerebellar input layer, granule cells, while larvae performed the closed-loop paradigm.

We first characterized the response types present in the granule cell population. Regression-based clustering (see Methods) of ROIs using sensory and motor events revealed functional clusters representing stimulus and turn-related activity in every larva (Figure 4a, Figure S4a, n=20), indicating that granule cells represent both the motor actions and the sensory context in which they are executed [40, 41].

**Figure 4:**
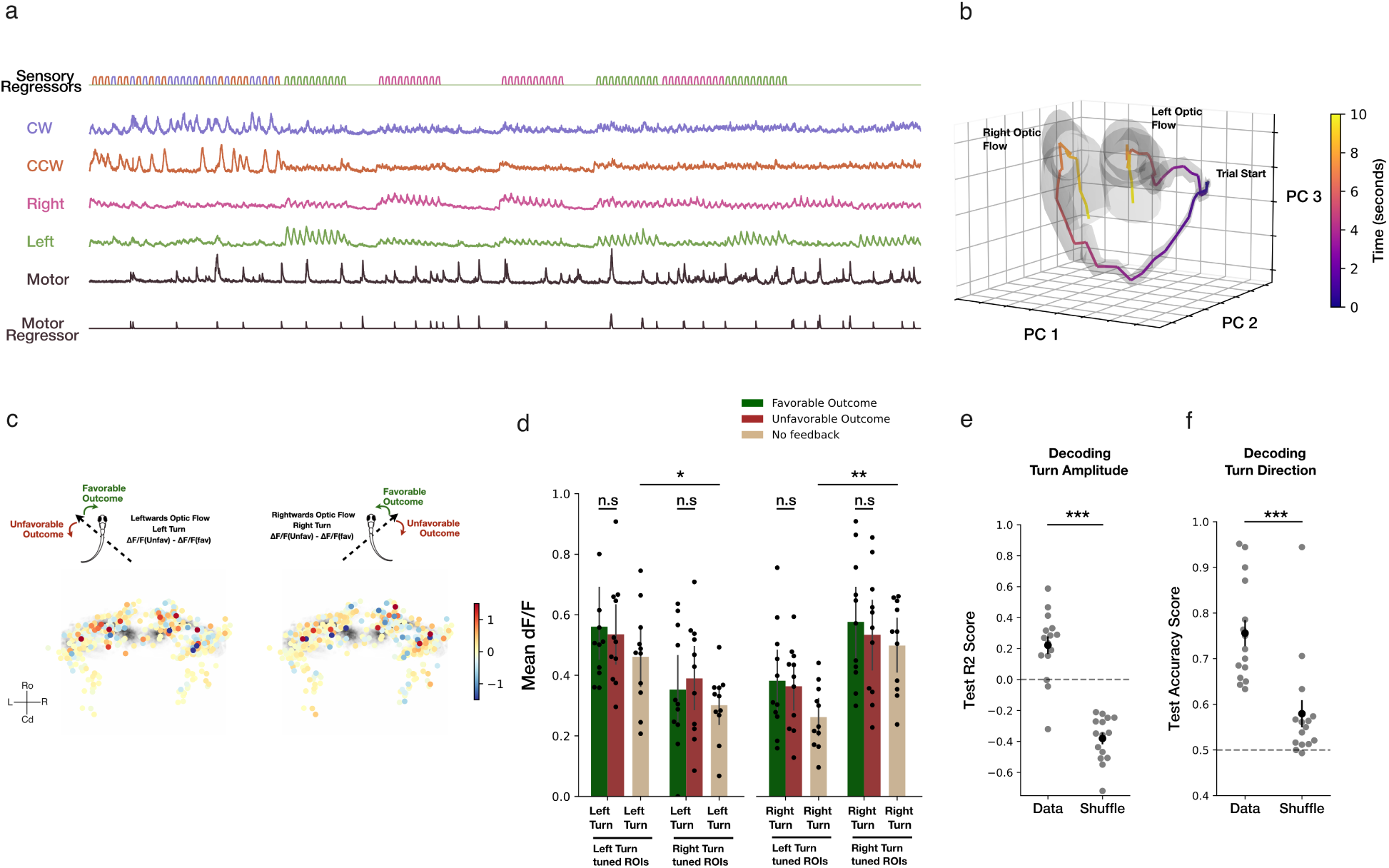
Granule cells provide an outcome-invariant sensorimotor basis. **a.** Functional clustering of granule cell activity in an example larva. ROIs are grouped by correlation with sensory and motor regressors, revealing distinct sensory- and turn-related clusters. **b.** Sensory-related granule cell activity projected into principal-component space. Rightward and leftward optic flow occupy distinct regions, showing spatiotemporal separation of the two stimuli. **c.** Anatomical map of motor-tuned granule cells (top 10-percentile of correlation with the motor regressor, pooled across larvae), coloured by the difference in activity between unfavorable and favorable outcome conditions (unfav − fav) following left turns during leftward flow (left) and right turns during rightward flow (right). **d.** Mean ΔF/F following left and right turns with favorable and unfavorable outcome, and no-feedback conditions, for left- and right-tuned ROIs (each point, one larva). **e.** Cross-validated decoding of turn amplitude from motor-related granule cell activity **f.** Cross-validated decoding of turn direction from motor-related granule cell activity. All plots show mean and sem (shaded areas and error bars) across fish (n=16 fish).

We observed that the sensory responses were low-dimensional but structured. Principal component analysis of sensory ROIs (see Methods) showed that a median of 4 principal components (interquartile range, IQR=2.5) accounted for 90% of the variance (Figure S4d right). Despite this low dimensionality, rightward and leftward optic flow exhibit distinct spatiotemporal patterns (Figure 4b, Figure S4d left) in neuronal activity space, and individual ROIs are active at distinct time points within a trial (see Methods), tiling the stimulus period [42] (Figure S4f,g). Thus, granule cell activity forms a temporally organized basis that encodes both stimulus identity and elapsed time, thereby providing a structured representation for downstream cerebellar computations.

Next, we classified motor ROIs into left and right populations (top 10-percentile of correlations with left and right turns, respectively - see Methods) and compared their responses following turns generated by the innate policy, i.e., right turns during rightward optic flow and left turns during leftward optic flow. As expected, these ROIs show strong turn-direction specificity (Figure 4d) (left-ROIs leftturnΔf/f = 0.46 ± 0.04 vs rightturnΔf/f = 0.26 ± 0.03, Wilcoxon signed-rank test p < 0.01 and right-ROIs rightturn Δf/f = 0.49 ± 0.05 vs leftturn Δf/f = 0.30 ± 0.03, Wilcoxon signed-rank test p < 0.001). Notably, their responses did not differ by the outcome these turns produced under normal and inverse rule trials (Figure 4c,d). Using post-turn granule cell population activity, a multiclass logistic regression model with five-fold cross-validation (see Methods) did not decode the turn outcome significantly better than the label-shuffled distribution (Figure S4e) (33.63 ± 3.19% vs shuffle 32.31 ± 2.38% Wilcoxon signed-rank test p=0.44). This suggests that the turn outcome was not represented in a sufficiently distinct population activity pattern to support reliable decoding.

Despite this lack of outcome-dependent modulation, granule cell post-turn activity supported reliable decoding(see Methods) of motor kinematic parameters. Turn amplitude was predicted by multivariate linear regression (Figure4e, FigureS4i) (5-fold cross-validation test R-squared: data= 0.22 ± 0.05 vs shuffle= -0.38 ± 0.03, Wilcoxon signed-rank test p < 0.001), and turn direction by logistic regression (Figure 4f, Figure S4h) (5-fold cross-validation test accuracy: data= 75.5 ± 2.8% vs shuffle = 58 ± 2.9%, Wilcoxon signed-rank test p < 0.001). Granule cells, thus, carry reliable graded information about an animal’s own movement that is unaffected by the outcome of the movement.

Together, these results suggest that granule cell activity has a temporally structured representation of sensory context and motor actions, but not of the sensory consequences produced by those motor actions, indicating that granule cell activity provides an outcome-invariant sensorimotor basis for downstream computations [10, 43].

### Inferior olive neurons encode the sensory consequences of self-generated movements

Having established that granule cells encode contextual and motor information but not the sensory consequences of movements (Figure 4), we next examined the inferior olivary neurons (IONs), which provide the climbing fiber inputs to the Purkinje cells, during the same behavioral paradigm.

IONs show robust sensory responses to optic flow [44, 45] (Figure S5a) and also exhibit a post-turn increase in activity (Figure S5b), consistent with a signal driven by the sensory feedback that the movement generates. To test whether this signal is modulated by the sensory outcome of turns, we classified IO neurons into left and right populations based on anatomical location and examined their responses to turns generated by the innate policy. Unlike granule cells, ION activity was strongly modulated by the sensory outcome (Figure 5a). Left IONs responded more strongly to right turns with unfavorable outcome than with favorable outcome (unfav Δf/f = 0.12 ± 0.02 vs fav Δf/f = 0.05 ± 0.02, Wilcoxon signed-rank test with Benjamini-Hochberg correction for multiple comparisons p <0.05), while responses to left turns were similar regardless of the outcome (p=0.617). Right IONs showed a mirrored response pattern such that they responded more to left turns with unfavorable outcome than with favorable outcome (unfav Δf/f = 0.14 ± 0.04 vs favΔf/f = 0.09 ± 0.01, Wilcoxon signed-rank test with Benjamini-Hochberg correction for multiple comparisons p <0.05), while responses to right turns were unchanged (p = 0.617). IONs thus exhibit selectively enhanced activity when an action produced an unexpected sensory outcome (Figure 5c).

**Figure 5:**
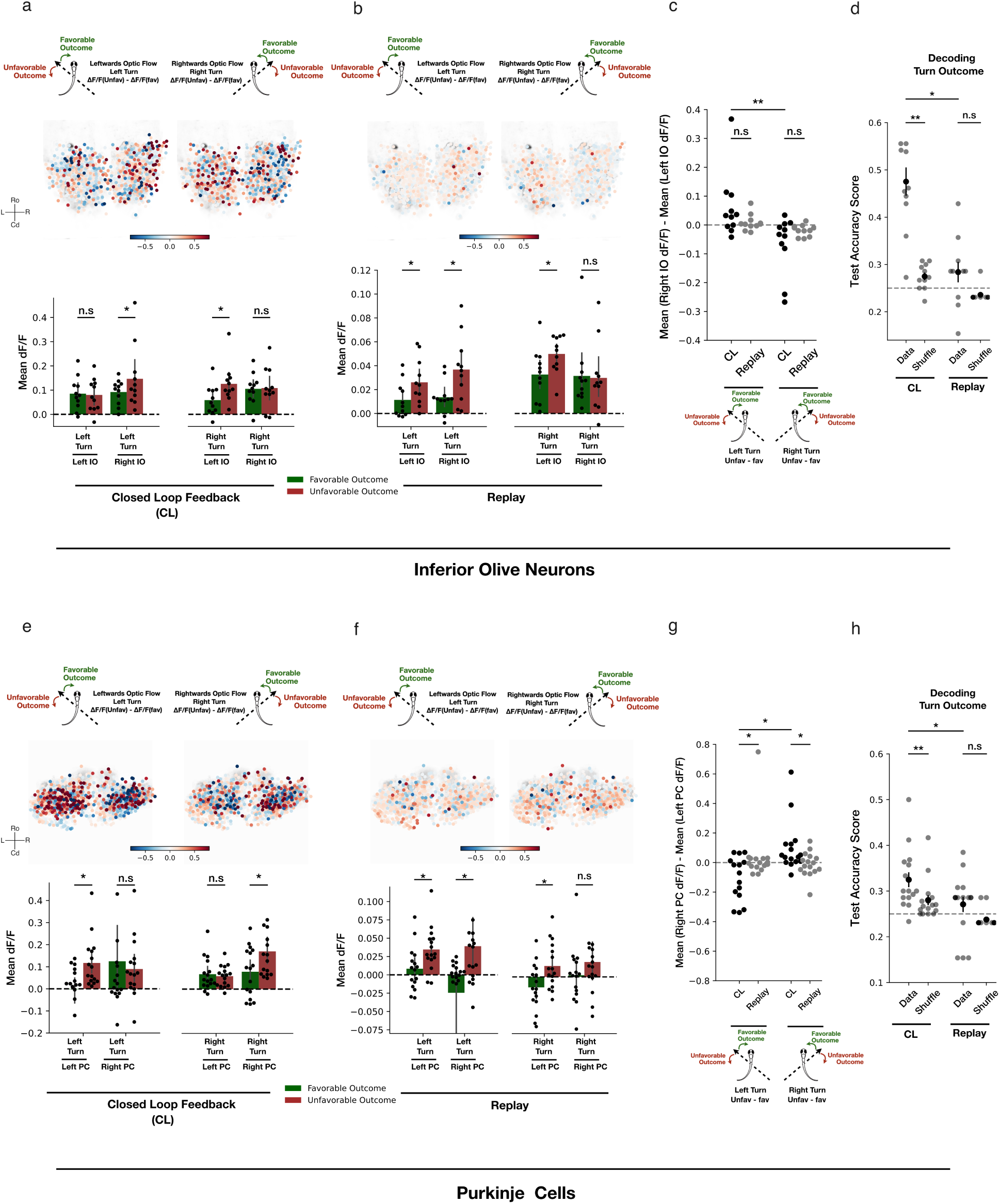
Outcome-dependent modulation of inferior olive neurons and Purkinje cells. **a.** Anatomical map of inferior olive neurons (ROIs in the top-10 percentile correlation with sensory regressors across all fish) showing Unfav-fav activity for left turns during left optic flow (left) and right turns during right optic flow (right) during closed-loop behavior(top). Quantification of Mean df/f after left and right turns with favorable and unfavorable outcome conditions, split across left and right ION ROIs for all fish during closed-loop behavior(bottom). **b.** Anatomical map of inferior olive neurons (ROIs in the top-10 percentile correlation with sensory regressors across all fish) showing Unfav-fav activity for left turns during left optic flow (left) and right turns during right optic flow (right) during replay(top). Quantification of Mean df/f after left and right turns with favorable and unfavorable outcome conditions, split across left and right ION ROIs for all fish during replay(bottom). **c.** Mean difference in activity between right- and left-tuned inferior olive ROIs during closed-loop behavior and replay conditions, quantified separately for left- and right-Unfav-fav differences. **d.** Decoding performance of a multiclass logistic regression model (5-fold cross-validation) trained to jointly predict turn outcome from inferior olive population activity during closed-loop behavior and replay. All plots show mean and sem (error bars) across fish (n=11 ION fish). **e.** Anatomical map of Purkinje cells (ROIs in the top-10 percentile correlation with sensory regressors across all fish) showing Unfav-fav activity for left turns during left optic flow (left) and right turns during right optic flow (right) during closed-loop behavior(top). Quantification of Mean df/f after left and right turns with favorable and unfavorable outcome conditions, split across left and right Purkinje cell ROIs for all fish during closed-loop behavior(bottom). **f.** Anatomical map of Purkinje cells (ROIs in the top-10 percentile correlation with sensory regressors across all fish) showing Unfav-fav activity for left turns during left optic flow (left) and right turns during right optic flow (right) during replay(top). Quantification of Mean df/f after left and right turns with favorable and unfavorable outcome conditions, split across left and right Purkinje cell ROIs for all fish during replay(bottom). **g.** Mean difference in activity between right- and left-tuned Purkinje cell ROIs during closed-loop behavior and replay conditions, quantified separately for left- and right-Unfav-fav differences. **h.** Decoding performance of a multiclass logistic regression model (5-fold cross-validation) trained to jointly predict turn outcome from Purkinje cell population activity during closed-loop behavior and replay. All plots show mean and sem (error bars) across fish (n=16 PC fish).

Using ION post-turn activity, we could jointly decode turn outcome (Figure 5d) with 47.53 ± 2.9% (p < 0.01) test accuracy using a multiclass logistic regression model with 5-fold cross-validation (shuffle accuracy = 27.48 ± 0.8%, chance level=25%), confirming that ION activity carries structured information about the sensory consequences of motor actions.

However, a key question is whether this outcome dependence reflects the sensory consequences of the movement or simply the change in the visual stimulus itself. These are difficult to dissociate during closed-loop behavior as the animal’s turn and the resulting visual flow are coupled. To address this, we therefore performed replay experiments in which visual feedback generated during closed-loop behavior was replayed to the larva in an open loop. Replay stimuli were assigned a turn and outcome label corresponding to the behavioral events that generated them during closed-loop conditions (Figure S5g). If the inferior olive signals were purely sensory, replay should reproduce the closed-loop responses.

Instead, replay elicited only a partial recapitulation of the closed-loop activity. Replay evoked similar qualitative patterns (Figure 5b,c). However, the replay activity was half that seen during closed-loop behavior, and unlike closed-loop activity, turn outcome could not be decoded above chance levels (25%) from the replay responses (Figure 5d) (replay accuracy = 28.37 ± 2.1%, shuffle accuracy = 23.57 ± 0.4%, Wilcoxon signed-rank test p = 0.1). Although visual input contributes to inferior olive activity, the structured feedback-dependent modulation of ION activity requires the visual change to be a consequence of a self-generated movement.

Together, these results show that the IONs encode the sensory consequences of self-generated movements rather than visual stimuli alone. Their responses are selectively enhanced when movements produce unexpected sensory outcomes [46]. Such a representation is consistent with a sensory prediction-error signal that could provide the instructive input [47–50] required for Purkinje cell-dependent correction.

### Outcome-dependent Purkinje cell activity is positioned to drive corrective behavior

Having found motor and contextual representations in granule cells (Figure 4) and reafferent error signals in inferior olive neurons (Figure 5 top), we asked how these signals are represented at the level of Purkinje cells. To address this, we performed 2-photon imaging of Purkinje cells during the assay.

Purkinje cell activity is spatially organized and lateralized [51–53] such that clockwise rotations and rightward optic flow preferentially activated the left hemisphere (Figure S6a,d). In contrast, counterclockwise rotations and leftward optic flow activated the right hemisphere (Figure S6a,d). This lateralization was opposite to that observed in the inferior olive neurons (Figure S5e), consistent with the contralateral olivo-cerebellar projections [44].

We classified Purkinje cells into left and right populations based on anatomical location and compared their responses following turns generated by the innate policy. We observed that the Purkinje cell activity was also strongly modulated by the sensory outcome of the turns (Figure 5e). Left Purkinje cells responded more strongly following left turns with unfavorable outcome than favorable outcome (invΔf/f = 0.11 ± 0.02 vs normΔf/f = -0.001 ± 0.02, Wilcoxon signed-rank test with Benjamini-Hochberg correction for multiple comparisons p < 0.05), while responses to right turns remained unchanged. Similarly, right Purkinje cells responded more strongly following right turns with unfavorable outcome than favorable outcome (invΔf/f = 0.17 ± 0.03 vs normΔf/f = 0.07 ± 0.02, Wilcoxon signed-rank test with Benjamini-Hochberg correction for multiple comparisons p < 0.05), and responses to left turns remained unchanged. Thus, Purkinje cell responses were selectively enhanced in the hemisphere ipsilateral to the turn that produced an unfavorable outcome.

Just as we had seen for IONs, this modulation also required a self-generated movement. During replay, the same qualitative pattern appeared, but the responses were of reduced magnitude and lacked the spatial organization seen during closed-loop behavior (Figure 5f,g). Consistent with this, turn outcome could be jointly decoded (Figure 5h, see Methods) from Purkinje cell activity during closed-loop behavior (multiclass logistic regression with 5-fold cross-validation test accuracy = 32.47 ± 1.6 % vs shuffle accuracy = 27.4 ± 1.1% Wilcoxon signed-rank test p < 0.01) but not from the replay responses (test accuracy = 27.09 ± 2.3% vs shuffle accuracy = 23.76 ± 0.4% Wilcoxon signed-rank test p = 0.08).

Together with the loss of correction after Purkinje cell ablations observed above (Figure 3), these results indicate that Purkinje cells convert the sensory outcome of a movement into a signal that drives corrective behavior [54, 55]. We observed an increase in the Purkinje cell activity in the hemisphere ipsilateral to the turn that produced an unfavorable outcome. As Purkinje cells inhibit downstream circuitry through eurydendroid cells [56, 57], we propose that this enhanced activity transiently suppresses the action that produced the unfavorable outcome [58, 59] (Figure S6f), biasing the animal toward alternative turns with more favorable consequences. Such a mechanism would allow the reafferent error carried by the inferior olive to be converted, at the Purkinje cell, into the rapid trial-by-trial correction observed behaviorally.

### Innate feed-forward policy and cerebellar correction jointly determine adaptive behavior

We observe that the behavioral correction progressively weakens as the optic flow angle increases (Figure 2), raising the question whether this reflects a loss of cerebellar correction or a limitation imposed elsewhere in the circuit. To distinguish between these possibilities, we imaged inferior olive neurons and Purkinje cells with optic flow at 45° and 60° under both normal and inverse rules. Despite reduced behavioral correction at 60°, both IONs and Purkinje cells exhibited robust outcome-dependent activity, and the population responses reliably decoded turn outcome (Figure 6e,f) using a multiclass logistic regression classifier (see Methods). Thus, reduced behavioral correction at larger perturbations cannot be explained by a loss of the olivo-cerebellar error signal.

**Figure 6:**
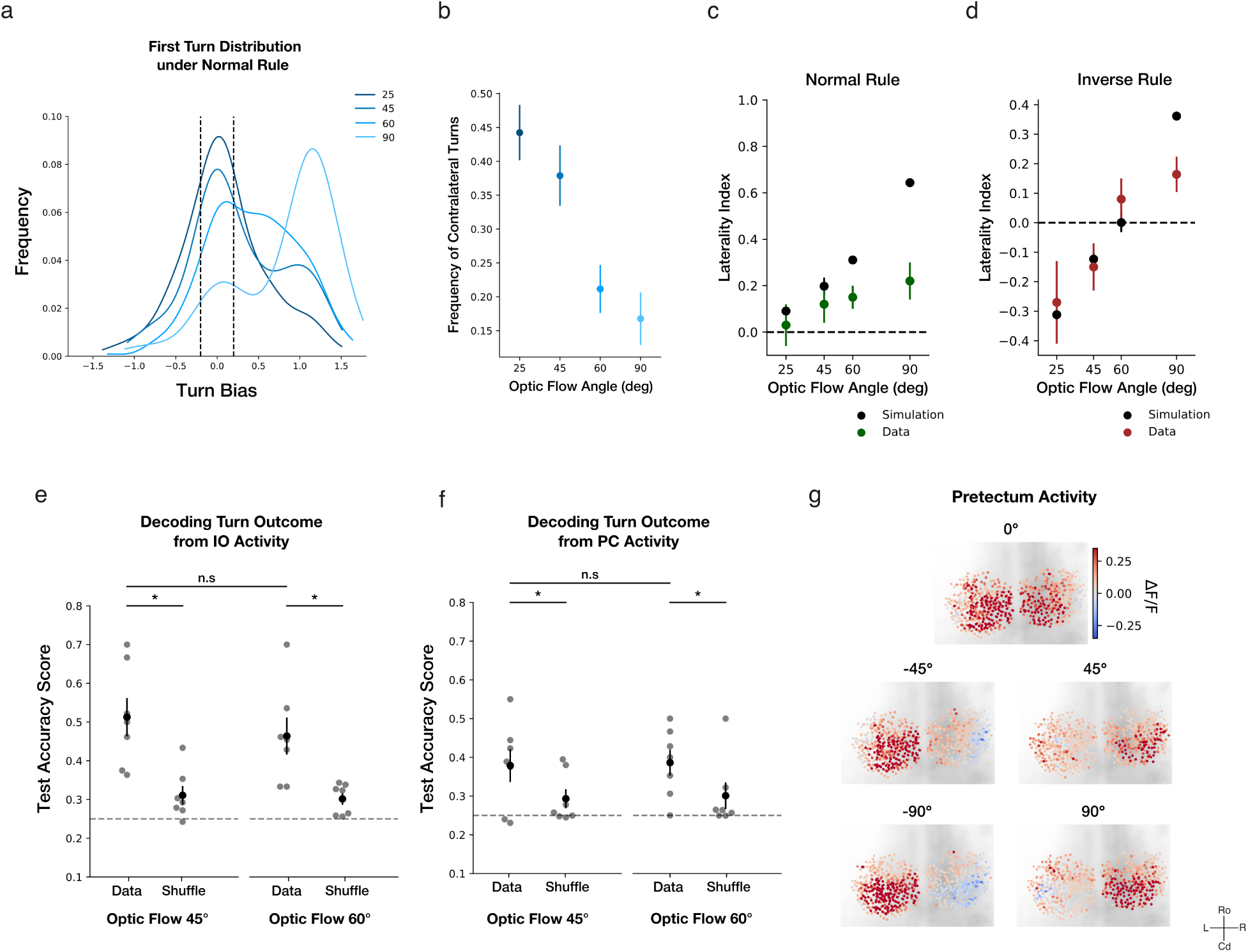
An innate feed-forward policy explains the limits of adaptive behavior. **a.** Distribution of the first turn angle chosen under normal rule across optic flow angles (25°, 45°, 60°, 90°). **b.** Mean frequency of contralateral turns under normal rule across optic flow angles. **c,d.** Laterality index across trials under normal rule and inverse rule for the model initialized with the measured innate policy and for larvae **e.** Decoding of turn outcome from inferior olive population activity (multiclass logistic regression, 5-fold cross-validation) during closed-loop behavior at 45° and 60°. **f.** Decoding of turn outcome from Purkinje cell population activity (multiclass logistic regression, 5-fold cross-validation) during closed-loop behavior at 45° and 60°. **g.** Pretectal ΔF/F maps for different directions of optic flow in a representative larva. All plots show mean and sem (shaded areas and error bars) across fish (n=18 fish for normal rule turn distribution, 7 fish for ION activity at 45° and 60°, 7 fish for PC activity at 45° and 60°, and 13 fish for pretectum activity). The pretectum data were reanalyzed from [62].

Instead, we observed that the innate stimulus-action mapping was not fixed and varied systematically with the optic flow angle (Figure 6a). Under normal rule, optic flow at 25° had a broad distribution of turn angles, with 44.23 ± 4.08% in the contralateral direction. As the angle increased, the turn distribution shifted towards more ipsilateral turns, and the fraction of contralateral turns dropped to 16.76 ± 3.89% at 90° (Figure 6b).

The zebrafish pretectum is a region that encodes translational whole-field motion and is believed to represent behavioral drive [16, 60, 61]. To check for any signature of this angle-dependent innate policy, we analyzed pretectal activity in a previously published whole-brain imaging dataset in which larvae viewed optic flow at 0°,± 45°,± 90°[62]. This revealed progressively stronger lateralization in the pretectal activity as the optic flow angle increased. At 0°, the left and right pretectum were equally activated. At larger optic flow angles, the activity progressively lateralized to the hemisphere ipsilateral to the flow (Figure 6g, Figure S7d,e). This pattern closely paralleled the behavioral policy such that optic flow angles that strongly elicited lateralized pretectal activity also generated predominantly ipsilateral turning responses and exhibited the weakest adaptive correction under inverse rule.

The behavioral and imaging results further suggest that the limits of correction are determined by the interaction between the instructive signal and the actions available within the innate policy. At smaller optic flow angles, the innate behavioral policy produces a greater proportion of contralateral turns, which under inverse rule leads to more favorable sensory outcomes. This suggests that actions already present in the innate repertoire that happen to produce a favorable outcome are being reinforced during inverse rule. Thus, behavioral correction is more effective at smaller angles because corrective contralateral turns are sampled more frequently and can therefore be preferentially reinforced. However, at larger angles, reduced sampling of corrective contralateral turns limits behavioral correction.

To test whether this structure of the innate behavioral policy is sufficient to explain the limits of behavioral correction, we developed an actor-critic reinforcement-learning simulation [8, 63, 64] (Figure S7a; see Methods). The agent selected a turn from the experimentally observed action repertoire for that particular optic flow angle, evaluated the sensory consequences of that action, and used the resulting error signal [48, 65] to update the action values, biasing future action selection. Importantly, the policy was initialized using the experimentally measured innate turn-angle distribution for each optic-flow angle during normal rule (Fig.6a), and all other parameters, such as the learning rates, were held constant across conditions.

We show that initializing the model with turn-angle distributions measured under the normal rule is sufficient to reproduce the pattern of behavioral correction observed across optic-flow angles. The model shows the strongest correction at 25°, which then progressively weakens with increasing optic flow angle, closely matching the experimental data (correlation=0.92) (Figure 6d), while the model correctly predicts the absence of adaptation under normal rule (correlation=0.98) (Figure 6c). In contrast, initializing the model with a uniform action policy eliminated this angle dependence of correction, producing equally strong correction across all optic flow angles (Figure S7b). These results demonstrate that the experimentally measured innate behavioral policy, rather than the learning rule itself, determines the limits of behavioral correction.

Together, these results suggest that this adaptive behavior emerges from the interaction of two complementary factors. An angle-dependent feed-forward sensorimotor policy, plausibly specified by the pretectum, that determines the distribution of actions available to the animal, whereas the olivo-cerebellar system evaluates the sensory consequences of those actions and biases future behavior. The error signal can only reinforce actions that are actually expressed [66, 67], so the effectiveness of cerebellar learning is constrained by the innate behavioral policy generated by the feed-forward pathway. As optic flow angle increases, the pretectum increasingly biases behavior toward large ipsilateral turns, progressively reducing the probability of sampling the corrective contralateral action. Consequently, although the olivocerebellar error signal remains intact, it has progressively fewer opportunities to reinforce the behavior required to resolve the visuomotor mismatch. Thus, adaptive behavior is not only determined by the cerebellar correction but also by the structure of the innate stimulus-action policy.

## Discussion

Animals often need to balance the stability of innate behavior with the flexibility required to respond to changing environments. Here, we show how these demands can be met through an interaction between an innate feed-forward sensorimotor policy and the feedback about the consequences of the action. When the sensory consequences of innate visuomotor responses were reversed, larvae rapidly generated corrective turns without remapping the underlying innate sensorimotor policy. This behavioral correction required Purkinje cells and was accompanied by structured representations of sensory and motor context and action outcomes across the olivo-cerebellar system. However, this correction was not solely determined by the action-outcome signal. As the optic flow angle increased, the correction progressively weakened, even though the inferior olives and Purkinje cells reliably reported the sensory consequences of self-generated actions. In parallel, the innate behavioral repertoire became increasingly biased toward larger ipsilateral turns, with the corrective contralateral actions becoming less frequent. An actor-critic framework constrained by the experimentally measured innate behavioral policy reproduced the emergence and limits of this correction. Together, these findings show an interaction between feed-forward and feedback-driven control in which the innate policy determines which actions are available and how frequently they are expressed, whereas an olivocerebellar instructive signal biases those actions according to their consequences.

The interaction between these two processes is evident in the way larvae adapt to inverse rule. Rather than remapping their innate policy, larvae initially turn ipsilaterally according to their innate policy and then subsequently turn in the corrective contralateral direction. This provides a potential solution to the competing demands of robustness and flexibility. An innate response can remain available as a rapid default while feedback-dependent mechanisms recruit corrective actions when its consequences become unfavorable. In this view, cerebellar adaptation need not construct an entirely new sensorimotor transformation but can instead modify the expression of an existing policy while leaving its feed-forward generation intact. This interpretation is consistent with our finding that Purkinje cell ablation abolished adaptive correction while preserving the innate optomotor response. However, these results should be interpreted in the context of the timescale of our experiments. Larvae underwent only 15 inverse rule trials, sufficient to show rapid correction but potentially too short for a complete remapping of the innate policy. Therefore, our results do not imply that innate policies are immutable but rather show that substantial behavioral correction can occur without replacing the underlying policy. Whether long-term exposure to inverse feedback leads to a complete remapping [21], and whether this remapping requires the same circuitry, remains an open question.

The structure of the innate policy also varied with the stimulus context. As the optic flow angle increased, the innate turn distribution under normal rule became increasingly biased towards ipsilateral turns. Neural activity in the pretectum, a major feed-forward pathway that drives the optomotor response [15, 16], mirrored this angle-dependent change in behavioral policy. Pretectal activity became progressively more lateralized toward the hemisphere ipsilateral to the optic flow as the angle increased, with the strongest lateralization at 90°, where behavior was most strongly biased toward ipsilateral turns. This provides a neural correlate for the angle-dependent feed-forward policy and suggests that the pretectum shapes the initial set of actions available to the fish. Importantly, this shift in the policy was accompanied by a progressive weakening of adaptive correction under inverse rule, linking the structure of the innate policy to the extent of subsequent behavioral flexibility.

The olivo-cerebellar circuit provides the necessary computations required for feedback-driven correction. Granule cells represented both sensory and motor context[40], but these representations were largely outcome-invariant. However, granule cell activity reliably encoded behavioral variables such as turn direction and amplitude. Rather than directly signaling whether an action was favorable or unfavorable, granule cells therefore appear to provide a structured representation of the current sensory and motor state that can serve as a basis for downstream evaluation of movement outcomes. In contrast, inferior olive neurons were selectively modulated by sensory consequences of self-generated movements. Replay experiments showed that inferior olive activity could not be explained by visual stimulus alone but required a reafferent coupling between movement and its outcome. This places the ION signal in a distinct computational context. The outcome in our experiments is neither an externally delivered reward nor a passive sensory violation but a sensory consequence of the animal’s own movement. Thus, the ION signal links the action, its sensory consequence, and subsequent behavioral adjustment variables, which are often studied separately. This provides a useful bridge between classical supervised-learning theories of the cerebellum [3–5, 32, 68] and more recent observations that climbing fibers participate in reinforcement and expectation-related computations [9, 11, 69]. These signals then converge onto the Purkinje cells, which are required for converting this outcome-dependent signal into behavioral correction without disrupting the feed-forward optomotor response. However, these results do not show how the observed Purkinje cell activity alters the probability of the next action. Understanding this mechanism will require spatiotemporal manipulation of Purkinje cell activity within our adaptive paradigm.

The interaction between a plausible pretectal feed-forward control and cerebellum-mediated rapid feedback-driven correction is further emphasized by the angle dependence of behavior correction. As mentioned previously, behavioral correction progressively weakens with increasing optic flow angle, yet this decline was not accompanied by a loss of inferior olive signal. At larger angles (60°), both ION and Purkinje cell populations continued to show structured outcome-dependent modulation and reliably encoded turn outcome. Thus, failure of correction could not be attributed to a loss of the olivo-cerebellar error signal. Instead, we see a strong shift in the innate policy as the optic flow angle increased, such that the turn distribution became increasingly concentrated on ipsilateral turns, while the corrective contralateral actions were sampled less frequently. This suggests that the effectiveness of cerebellar learning is constrained by the innate behavioral policy generated by the feed-forward pathway.

The actor-critic model provides a quantitative test for this interpretation. When initialized with experimentally measured turn distributions at different optic flow angles, the model reproduced the decline in correction with increasing optic flow angles without changing any learning parameters across conditions. In contrast, replacing the measured policy with a uniform policy eliminated angle dependence and yielded comparable corrections across all optic flow angles. Thus, the structure of the innate policy was sufficient to determine the capacity for subsequent adaptation. This interpretation gives a functional role to the variability of innate behavior. The distribution of turn directions is not simply noise around a fixed motor command [70], rather it determines which alternative actions the animal samples and, therefore, which action-outcome relationships become available for learning [66, 71]. Therefore, motor variability can provide a substrate for exploration rather than merely representing imprecision in motor control.

Together, these results show that feed-forward circuits provide a structured repertoire of actions and determine which behaviors are sampled, while feedback about the consequences of those actions modulates subsequent action selection. Thus, adaptive behavior is determined not only by the fidelity of the instructive signal but also by the structure of the behavioral repertoire on which that signal acts.

## Methods

### Zebrafish husbandry

All experiments involving larval zebrafish were performed in accordance with institutional and governmental regulations, with approvals obtained from the Technical University of Munich (TUM) and the regional authority (Regierung von Oberbayern).

Adult zebrafish were housed in a fish facility with a recirculating aquarium system with continuous carbon, biological, and UV filtration. The system included an automatic daily exchange of 12% water volume to ensure stable water quality. Environmental parameters were carefully controlled, with water temperature maintained between 27.5 °C and 28 °C, pH adjusted to 7.0–7.5, and conductivity was kept between 750-800 μS. Fish were maintained under a 14-10 hr light-dark cycle. Adults were housed in 3.5 L tanks at a density of 10–17 animals per tank. They were fed twice daily with a combination of Gemma Micron 300 (Skretting) and freshly hatched Artemia salina to provide a balanced diet.

All experiments were performed with zebrafish larvae aged 6–8 days post-fertilization (dpf). Breeding was carried out by placing 2–3 pairs of adult zebrafish together in a single spawning tank. Fertilized eggs were collected shortly after spawning and placed in 90 mm Petri dishes containing 0.3× Danieau’s solution, which prevented microbial contamination during the early developmental stages. Embryos were kept in this medium until 3 dpf, after which they were transferred into filtered fish facility water. The larvae were fed with rotifers before any experiments. Since zebrafish do not begin sexual differentiation until 20–25 dpf, the sex of the larvae used in this study is undetermined. Larvae used for imaging and ablation experiments were anesthetized using ice water at 4 or 5 dpf and screened for fluorescence under an epifluorescent microscope.

### Fish lines

The Tuepfel long-fin (TL) wild-type strain was used for head-restricted behavioral experiments. For imaging experiments, larvae with mitfa -/- mutation (nacre) were used. The following lines were used for experiments:

- Tg(PC:Ntr-tagRFP) [21] was used for Purkinje cell chemogenetic ablation.
- Tg(GR90:Gal4);Tg(UAS:GCaMP6s) [72] was used 2p-imaging of granule cells
- Tg(vglut2a:Gal4);Tg(UAS:GCaMP6s) [73] was used 2p-imaging of inferior olive neurons
- Tg(PC:GCaMP6s) [51] was used for 2p-imaging of Purkinje cells

### Head-restricted behavioral setup

Behavioral experiments were carried out in a custom-built setup [31]. Head-restricted larval zebrafish was positioned on an acrylic sheet with a screen mounted below it. Visual stimulus was projected onto the screen using an Asus P3E projector via a cold mirror (Edmund Optics) placed at 45°below the screen. An array of infrared (IR) LEDs was used to illuminate the fish from below, enabling tail tracking without interfering with visual stimulation. The camera was equipped with an objective lens (Navitar) for magnification and an IR-pass filter (Edmund Optics) to exclude visible light from the projected stimuli. The entire setup was housed in a dark aluminum cage to eliminate any interference from external light sources. Fish behavior (tail movements) was recorded at 200 Hz using a high-speed camera (Ximea).

### 2p-Imaging

Functional imaging experiments were performed using a custom-built 2-photon imaging system [62, 74–76]. A Ti:Sapphire laser (Spectra-Physics Mai Tai) tuned to 905 nm, with an approximate power of 10 mW at the back aperture of the imaging objective, was used for imaging. All fish were imaged at 3 frames per second. Only 1 plane was imaged per fish. The microscope was controlled using custom-made software (Brunoise, https://github.com/portugueslab/brunoise).

### Fish Preparation

Larval zebrafish were embedded using 2% low-melting-point agarose (ThermoFisher Scientific) in a 35mm Petri dish (Corning). The agarose surrounding the tail was carefully removed with scalpels, allowing the tail to move freely. The agarose surrounding the eyes was not removed, hence there was minimal to no eye movement during these experiments. The embedded larvae were allowed to recover for at least 2 hours before being transferred to the behavioral/imaging setup.

### Behavior Stimulus Protocols

For both behavioral and imaging experiments, larvae were presented with a pink noise pattern moving at 25 °, 45 °, 60 °, or 90 ° to the right or left, with a normal (norm) or inverse (inv) rule. The experiments consisted of 4 blocks (rightward motion with normal rule, leftward motion with normal rule, leftward motion with inverse rule, rightward motion with inverse rule), with 10-15 trials each with 15 s trial duration and 10 s inter-trial interval.

For imaging experiments, 3 additional blocks were presented: rotational block, no-feedback block, and replay block. In the rotational block, the pink noise pattern rotated either clockwise or counterclockwise for 15 seconds, followed by a 10-second pause. Each rotational direction was presented 10 times in a randomized order. The no-feedback block consisted of a pink noise pattern moving either to the right or left of the fish without any visual feedback. For the replay block, we recorded the visual stimuli as the fish interacted with the four experimental blocks during behavioral experiments and subsequently presented this recorded pattern to the fish in an open-loop format.

Stimulus presentation, real-time tracking, and data acquisition were controlled by a custom Python script written in Stytra [31]. The software was calibrated before the experiments to ensure accurate coordinate transformations between the digital and real world.

### Closed-loop design

The Stytra algorithm [31] tracks tail movements by fitting 9-16 line segments to the tail and computing the tail angle as the difference between the first and last segments. This tail angle, binned every 70ms and multiplied by a decaying exponential function (2.5 *∗ e^−^*^6.5*t*^)(t is the time measured from the start of the turn until the turn ends), is used to update the angle of the visual stimuli in real-time. This ensures that only the initial 150ms of a turn causes the maximum change in the stimulus angle (Figure S1a).

### Behavioral Data Analysis

For each fish, experiments were conducted in both normal (norm) and inverted (inv) rule, with optic flow moving either to the right (+45°) or to the left (−45°). To facilitate comparison across trials and animals, turning responses recorded during leftward optic flow trials were sign-inverted. Consequently, all trials were represented in a common reference frame in which the direction of optic flow was treated identically. Turns were therefore classified as either ipsilateral (occurring in the same direction as the optic flow) or contralateral (occurring in the opposite direction to the optic flow).

Turning events(bouts) were detected using a lab written software [77]. Turning behavior was quantified using the turn bias and turn vigor. Turn bias refers to the amplitude of tail deflection whereas turn vigor is defined as the standard deviation of the tail traces in the 70ms window after turn onset. To focus on turns associated with clear directional decisions, only turns with an absolute bias value greater than 0.2 were included in the analysis. Turns with a bias greater than 0.2 were classified as ipsilateral turns, whereas turns with a bias less than −0.2 were classified as contralateral turns. Turns with bias values between −0.2 and 0.2 were excluded from further analysis. Turns were also classified based on the outcome such that turns that rotated the optic flow toward the fish were classified as favorable turns and turns that rotated the optic flow away from the fish were classified as unfavorable turns.

### Laterality Index

To quantify the turning preference of the fish during each trial, we computed a Laterality index (LI) defined as:

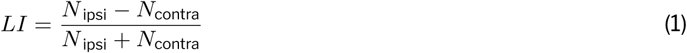

where *N* _ipsi_ is the number of ipsilateral turns and *N*_contra_ is the number of contralateral turns.

LI is bounded between *−*1 and +1. A value of +1 indicates that all classified turns were ipsilateral, whereas a value of *−*1 indicates that all classified turns were contralateral. An LI of 0 reflects no directional preference, corresponding to equal numbers of ipsilateral and contralateral turns. Trials with no turns were removed in this analysis.

### Chemogenetic Ablations

A nitroreductase (Ntr)-Nifurpirinol (NfP) chemogenetic approach [26, 78] was used to achieve targeted neuronal ablation. To target Ntr expression in zebrafish, we used the PC:Ntr-tagRFP transgenic line, which drives expression specifically in Purkinje cells. Prior to ablation, larvae were screened for tagRFP expression in Purkinje cells using an upright epifluorescence microscope (Leica). RFP-positive larvae were transferred to 90 mm Petri dishes with minimal water, and 30 ml of a 5 μM NfP solution in 0.2% DMSO was added to each dish.

The Petri dishes containing larvae were incubated at 28 °C for 18 hours in a dark box to prevent degradation of the photosensitive Nfp. After incubation, larvae were immediately removed from the NfP solution and washed three times with fresh fish water to ensure complete removal of NfP and DMSO. This method enables highly specific, reproducible ablation of neurons, allowing the functional consequences of their loss to be assessed in vivo [21, 62, 76, 79, 80].

### Regression-based Clustering

To identify functional clusters in granule cell, Purkinje cell, and inferior olive populations, we first constructed regressors for both sensory and motor events that happened during the experiment. These included clockwise and counterclockwise rotation, rightward and leftward optic flow, and rightward and leftward turns. Each regressor was convolved with a GCaMP6s kernel with a time constant of 1.5 s to account for calcium indicator dynamics.

We then computed the Pearson correlation between each ROI and each of these convolved regressors. For each regressor, correlation values from all ROIs across all fish were pooled. A threshold was defined as the 90th percentile of this pooled correlation distribution. Subsequently, within each fish, ROIs whose correlation values exceeded the corresponding threshold were assigned to the functional cluster associated with that regressor.

### Granule cell PCA

To characterize the population dynamics of granule cell activity [81], we reduced the dimensionality of the neural responses using principal component analysis (PCA). First, rightward and leftward optic flow trials were separated. We used the mean activity of rightward optic flow trials as the reference trial and used this to define the PCA space. PCA was performed on this activity matrix, and principal components were retained until the cumulative explained variance reached 90%. The retained components defined the low-dimensional state space for subsequent analyses.

All remaining trials were subsequently projected onto this reference PCA space, yielding low-dimensional trajectories representing the temporal evolution of population activity. As the stimulus identity of each projected trajectory was known, trajectories were grouped according to their corresponding optic flow direction.

To quantify the consistency of trajectories within a stimulus condition and the separation between stimulus conditions, a mean trajectory was calculated for each group by averaging the projected trajectories within each stimulus class. For every individual trajectory, the cosine distance to the mean trajectory of its own group was computed as a measure of within-group variability (intracluster distance). In addition, the cosine distance between each trajectory and the mean trajectory of the opposite stimulus group was calculated as a measure of between-group separation (intercluster distance). All distance calculations were performed in the PCA space using the complete set of retained principal components required to explain 90% of the variance.

### Center of Mass Sorting

To characterize the temporal structure of granule cell sensory responses [42], we averaged activity across right-ward and leftward optic flow trials for all ROIs. Only ROIs with a Pearson correlation greater than the 85-percentile threshold of the population-wide correlation distribution with sensory regressors were included for this analysis.

For each ROI, the temporal activity profile was computed by calculating the cumulative integral of the mean response over time. The center of mass (COM) is defined as the time point at which the cumulative integral reached 50% of its total value. ROIs were sorted in ascending order of COM values to visualize the temporal ordering of activity across the population. We compare the distribution of COM values for granule cell populations with that for the inferior olive populations, showing a significantly narrower distribution for the latter.

### ΔF/F Computation

All functional imaging data were motion corrected, and regions of interest (ROIs) were extracted using suite2p [82]. For each ROI, baseline fluorescence(F0) was defined as the median over the entire experiment, and ΔF/F was computed as (F(t)-F0)/F0, where F(t) is the instantaneous fluorescence. To examine activity associated with an individual motor event (turn), ΔF/F traces were aligned to the onset of each bout. For each turn, ΔF/F trace was extracted from 0.5 sec before to 1 sec after the turn onset. This activity was analyzed separately by turn direction and outcome.

### Decoding Analysis

To determine whether granule cell population activity contained information about behavioral variables, we trained a linear regression model to predict turn amplitude and a logistic regression model to decode turn direction using the scikit-learn package [83]. For granule cell, Purkinje cell, and inferior olive populations, we trained a multiclass logistic regression model to decode the combined turn direction × outcome, with four classes corresponding to right turn-favorable outcome, right turn-unfavorable outcome, left turn-favorable outcome, and left turn-unfavorable outcome. For each turn, the neural activity of all ROIs within 1 s after turn onset was used as the feature vector.

Model performance was evaluated separately for each fish using five-fold stratified cross-validation, ensuring that the relative proportion of each class was preserved across training and test folds. Performance was then averaged across fish. To assess statistical significance, we generated a label-shuffled null distribution by randomly permuting the behavioral labels while preserving the neural activity. The decoding procedure was repeated for 100 shuffles to generate a null distribution. Comparisons across visuomotor rules, stimulus angles, and cell types used the same pre-processing, feature selection, and cross-validation procedures.

### Actor-Critic Simulation

To test whether the structure of the innate behavioral policy is sufficient to account for the angle-dependence of behavioral correction, we simulated this adaptation process using an actor-critic reinforcement learning framework [8] in which the only quantity that varied across optic flow angles was the innate behavioral policy measured during normal rule trials.

#### Action Space and Policy

The agent selected a turn from a discrete set of turn angles, with positive values indicating ipsilateral turns and negative values indicating contralateral turns, matching the convention used for behavioral data.

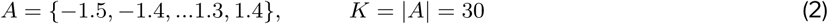

The actions were selected using a softmax policy over a vector of action preferences *w ∈* R*^K^*.

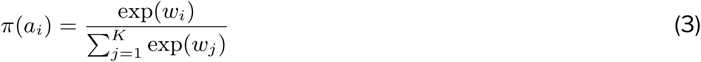

The action preferences that formed the agent’s initial policy were initialized using behaviorally measured turn distributions during normal rule trials. For a given angle *θ*, 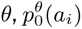 denotes the behaviorally measured, normalized distribution of turn angles obtained from the normal rule trials. We set

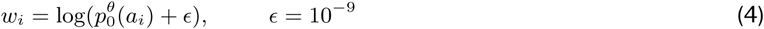

which gives 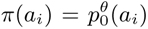 for the first action. As learning acts additively on *w* while behavior is read out through the softmax, this initialization seeds the learnable preferences with the animal’s innate action prior. The measured prior 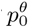 is the only model component that differed across optic flow angles.

#### Reward

The sensory consequences of each turn was scored using an angle-independent reward function.

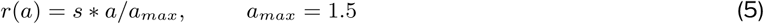

where *s* = *−*1 under inverse rule and *s* = +1 under normal rule, such that the corrective action (contralateral turns under inverse rule and ipsilateral turns under normal rule) is rewarded.

#### Teaching signal and learning

The learning was driven by a scalar reward-prediction error, which represents the inferior olive teaching signal [48]. A critic maintains a estimate of expected reward *V*, updated after each turn,

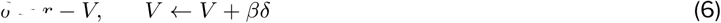

with critic learning rate *β* = 0.28. The action preferences were updated using a policy-gradient (REINFORCE) rule using this teaching signal,

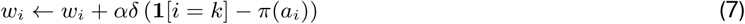

where *k* is the index of action taken, **1**[.] is the indicator function and *α* = 0.32 is the policy learning rate. Each simulation session consisted of 15 normal/inverse rule trials with 6 turns each, where the preferences *w* and critic *V* were updated at every turn and carried across trials within a session. For each trial, we computed the laterality index matching the behavioral definition. We simulated 100 agents per condition and reported the mean±s.e.m. across agents.

All model parameters (*α*, *β*, the action set *A*, and the numbers of bouts, trials and agents) were fixed across all visual stimuli angles and visuomotor rules; the only quantity that varied across optic flow angles was the measured innate prior 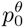. To assess whether the angle dependence of behavioral correction depended on specific model parameters, we systematically varied the policy learning rate and critic learning rate parameters. For each parameter combination, we quantified the root-mean-squared error (RMSE) between model and experimental values, showing that a low RMSE across a broad region of parameter space (Figure S7c).

We emphasize that the model is intended as a minimal, sufficiency-level account rather than an identification of the animal’s learning algorithm. It shows that a single teaching signal acting on the measured innate policy is sufficient to reproduce the graded dependence of correction on optic flow angle.

## Statistics

For all figures, statistical significance is indicated as follows:- *: p<0.05, **: p<0.01, ***: p<0.001, and ns: not significant. All statistical tests are two-sided unless otherwise indicated.

Figure 1e (top): Trialwise normal rule vs inverse rule laterality index, Wilcoxon signed-rank test with Benjamini-Hochberg correction for multiple comparisons. p-values as indicated in the plot.

Figure 1e (bottom): Latency of first contralateral turn at trial 1 6.64± 0.49 sec v/s trial2 4.62 ± 0.43 sec - p<0.001; trial1 6.64± 0.49 sec v/s trial3 4.16 ± 0.40 sec - p<0.01; trial1 6.64± 0.49 sec v/s trial15 4.37 ± 0.4 sec -p<0.01. Wilcoxon signed-rank test with Benjamini-Hochberg correction for multiple comparisons.

Figure S1b: Mean ipsilateral turn rate - Mean contralateral rate for normal rule 0.043 ± 0.015 Hz vs inverse rule -0.042 ± 0.023 Hz p<0.001 calculated within 4-13 sec time window, Wilcoxon signed-rank test.

Figure S1c (left): Mean optic flow direction during normal rule vs inverse rule trials calculated within the 5-15 sec time window, p <0.001, Wilcoxon signed-rank test.

Figure S1c (right): Mean optic flow direction for inverse rule (trial1-3) vs inverse rule (trial 4-15) calculated within the 5-15 sec time window p <0.001, Wilcoxon signed-rank test.

Figure S1e: Mean turn duration normal rule 0.57 ± 0.03 sec vs inverse rule 0.57 ± 0.04 sec p=0.35, Mean time to next turn normal rule 3.45 ± 0.22 sec vs inverse rule 3.79 ± 0.29 sec p=0.39, Mean turn vigor normal rule 1.16 ± 0.03 rad vs inverse rule 1.06 ± 0.03 rad p<0.001, Wilcoxon signed-rank test with Benjamini-Hochberg correction for multiple comparisons.

FigureS1g: Mean turn duration favorable outcome 0.42 ± 0.02 sec vs unfavorable outcome 0.49 ± 0.02 sec p<0.001, Mean time to next turn favorable outcome 2.44 ± .018 sec vs unfavorable outcome 2.63 ± 0.15 sec p<0.05, Mean turn vigor favorable outcome 1.15 ± 0.03 rad vs unfavorable outcome 1.16 ± 0.03 rad p=0.35, Wilcoxon signed-rank test with Benjamini-Hochberg correction for multiple comparisons.

Figure S2b: For normal rule trials, laterality index with angle Linear mixed-effects model, LI *∼* Rank(Angle) + (1|fish) slope = 0.06 ± 0.01, intercept = −0.02, p < 0.0001. For inverse rule trials, laterality index with angle Linear mixed-effects model, LI *∼* Rank(Angle) + (1|fish) slope = 0.156 ± 0.02, intercept = −0.44, p < 0.0001

Figure 3b (top): Mean ipsilateral turn rate - Mean contralateral rate for normal rule 0.06 ± 0.02 Hz vs inverse rule -0.04 ± 0.03 Hz p<0.01 calculated within 4-13 sec time window for control fish, Wilcoxon signed-rank test.

Figure 3b (bottom): Mean ipsilateral turn rate - Mean contralateral rate for normal rule 0.07 ± 0.02 Hz vs inverse rule 0.05 ± 0.02 Hz p=0.67 calculated within 4-13 sec time window for PC ablated fish, Wilcoxon signed-rank test.

Figure S3c: Trialwise normal rule vs inverse rule laterality index for control fish, Wilcoxon signed-rank test with Benjamini-Hochberg correction for multiple comparisons. p-values as indicated in the plot.

Figure S3d: Trialwise normal rule vs inverse rule laterality index for PC ablated fish, Wilcoxon signed-rank test with Benjamini-Hochberg correction for multiple comparisons. p-values as indicated in the plot.

Figure S3e: Mean optic flow direction for inverse feedback (control fish) vs inverse feedback (PC ablated fish) calculated within the 5-15 sec time window p <0.001, Wilcoxon signed-rank test.

Figure S3f: Mean turn rate during normal rule trials for control fish 0.29 ± 0.03 Hz vs ablated fish 0.29 ± 0.02 Hz p =0.58, Mean turn rate during inverse rule trials for control fish 0.33 ± 0.03 Hz vs ablated fish 0.34 ± 0.02 Hz p =0.58. Mann-Whitney U test

Figure 4d: Mean ΔF/F left turn favorable outcome 0.56 ± 0.06 vs unfavorable outcome 0.53 ± 0.05 p= 0.831 for left tuned ROIs. Mean ΔF/F right turn favorable outcome 0.57 ± 0.06 vs unfavorable outcome 0.53 ± 0.06, p = 0.519 for right-tuned ROIs. Mean ΔF/F left turn favorable outcome 0.35 ± 0.06 vs unfavorable outcome 0.38 ± 0.05 p = 0.519 for right tuned ROIs. Mean ΔF/F right turn favorable outcome 0.38 ± 0.05 vs unfavorable outcome 0.36 ± 0.04 p = 0.7 for left tuned ROIs. Mean ΔF/F left turn no feedback left tuned ROIs 0.46 ± 0.05 vs right tuned ROIs 0.26 ± 0.03 p<0.05 and Mean ΔF/F right turn no feedback left tuned ROIs 0.30 ± 0.03 vs right tuned ROIs 0.49 ± 0.04 p<0.05. Wilcoxon signed-rank test with Benjamini-Hochberg correction for multiple comparisons.

Figure 4e: Test R2 score 5-fold cross-validation for decoding turn amplitude data 0.22 ± 0.06 vs shuffle -0.38 ± 0.04 p<0.001, Wilcoxon signed-rank test.

Figure 4f: Test accuracy score 5-fold cross-validation for decoding turn direction data 0.75 ± 0.03 vs shuffle 0.58 ± 0.03 p<0.001, Wilcoxon signed-rank test.

Figure S4d: Mean cosine distance intra-cluster 0.32 ± 0.02 vs inter-cluster 0.93 ± 0.05 p<0.001, Wilcoxon signed-rank test.

Figure S4e(left): Test accuracy score 5-fold cross-validation for decoding turn-outcome data 0.33 ± 0.03 vs shuffle 0.32 ± 0.02 p=0.44, Wilcoxon signed-rank test.

Figure S4e (left): Test accuracy score 5-fold cross-validation for decoding turn-outcome data vs shuffle at 4 time points. -0.3 sec 0.37 ± 0.04 vs 0.31 ± 0.02 p= 0.23, 0.3 sec 0.36 ± 0.05 vs 0.31 ± 0.02 p = 0.24, 0.6 sec 0.36 ± 0.03 vs 0.31 ± 0.02 p=0.20, 1 sec 0.33 ± 0.03 vs 0.31 ± 0.02 p=0.24. Wilcoxon signed-rank test with Benjamini-Hochberg correction for multiple comparisons.

Figure S4g: Distribution of center of mass for granule cell ROIs vs inferior olive ROIs, Kolmogorov–Smirnov test p<0.001.

Figure S4h: Test accuracy score 5-fold cross-validation for decoding direction data vs shuffle at 4 time points. - 0.3 sec 0.61 ± 0.02 vs 0.59 ± 0.03 p = 0.06, 0.3 sec 0.65 ± 0.03 vs 0.58 ± 0.03 p<0.01, 0.6 sec 0.77 ± 0.03 vs 0.57 ± 0.03 p<0.01, 1 sec 0.76 ± 0.03 vs 0.58 ± 0.03 p<0.01. Wilcoxon signed-rank test with Benjamini-Hochberg correction for multiple comparisons.

Figure S4i: Test R2 score 5-fold cross-validation for decoding turn amplitude data vs shuffle at 4 time points. -0.3 sec -0.10 ± 0.04 vs -0.22 ± 0.02 p<0.01, 0.3 sec -0.009 ± 0.06 vs -0.41 ± 0.05 p<0.001, 0.6 sec 0.20 ± 0.06 vs - 0.38 ± 0.05 p<0.001, 1 sec 0.27 ± 0.05 vs -0.4 ± 0.07 p<0.001. Wilcoxon signed-rank test with Benjamini-Hochberg correction for multiple comparisons.

Figure 5a: Left IO left turn favorable outcome ΔF/F 0.08 ± 0.02 vs unfavorable outcome ΔF/F 0.08 ± 0.02 p=0.617, Right IO left turn favorable outcome ΔF/F 0.09 ± 0.01 vs unfavorable outcome ΔF/F 0.14 ± 0.04 p<0.05, Left IO right turn favorable outcome ΔF/F 0.05 ± 0.02 vs unfavorable outcome ΔF/F 0.12 ± 0.02 p<0.05, Right IO right turn favorable outcome ΔF/F 0.1 ± 0.02 vs unfavorable outcome ΔF/F 0.1 ± 0.02 p=0.617 during closed-loop behavior, one-sided Wilcoxon signed-rank test with Benjamini-Hochberg correction for multiple comparisons.

Figure 5b: Left IO left turn favorable outcome ΔF/F 0.01 ± 0.004 vs unfavorable outcome ΔF/F 0.02 ± 0.005 p<0.05, Right IO left turn favorable outcome ΔF/F 0.013 ± 0.004 vs unfavorable outcome ΔF/F 0.04 ± 0.008 p<0.05, Left IO right turn favorable outcome ΔF/F 0.03 ± 0.006 vs unfavorable outcome ΔF/F 0.05 ± 0.004 p<0.05, Right IO right turn favorable outcome ΔF/F 0.031 ± 0.008 vs unfavorable outcome ΔF/F 0.029 ± 0.008 p=0.55 during replay, one-sided Wilcoxon signed-rank test with Benjamini-Hochberg correction for multiple comparisons.

Figure 5c: Mean Right IO ΔF/F - Mean Left IO ΔF/F for left turn Unfav-fav for closed loop 0.06 ± 0.03 vs replay 0.008 ± 0.008 p=0.206, Mean Right IO ΔF/F - Mean Left IO ΔF/F for right turn Unfav-fav for closed loop -0.06 ± 0.03 vs replay -0.02 ± 0.006 p=0.206, Mean Right IO ΔF/F - Mean Left IO ΔF/F for left turn Unfav- fav 0.06 ± 0.03 vs right turn Unfav-fav -0.06 ± 0.03 p<0.01 during closed-loop. Wilcoxon signed-rank test with Benjamini-Hochberg correction for multiple comparisons.

Figure 5d: Test accuracy score 5-fold cross-validation for decoding turn-outcome data 47.53 ± 2.9% vs shuffle 27.48 ± 0.8% p<0.01 for closed loop. Test accuracy score 5-fold cross-validation for decoding turn-outcome data 28.37 ± 2.1% vs shuffle 23.57 ± 0.2% p=0.101 for replay. Test accuracy score 5-fold cross-validation for decoding turn-outcome closed-loop data 47.53 ± 2.9% vs replay data 28.37 ± 2.1% p<0.05. Wilcoxon signed-rank test with Benjamini-Hochberg correction for multiple comparisons.

Figure 5e: Left PC left turn favorable outcome ΔF/F -0.001 ± 0.02 vs unfavorable outcome ΔF/F 0.11 ± 0.02 p<0.05, Right PC left turn favorable outcome ΔF/F 0.12 ± 0.08 vs unfavorable outcome ΔF/F 0.08 ± 0.03 p=0.57, Left PC right turn favorable outcome ΔF/F 0.06 ± 0.01 vs unfavorable outcome ΔF/F 0.05 ± 0.01 p=0.7, Right PC right turn favorable outcome ΔF/F 0.07 ± 0.02 vs unfavorable outcome ΔF/F 0.17 ± 0.03 p<0.05 during closed-loop behavior, one-sided Wilcoxon signed-rank test with Benjamini-Hochberg correction for multiple comparisons.

Figure 5f: Left PC left turn favorable outcome ΔF/F 0.008 ± 0.007 vs unfavorable outcome ΔF/F 0.03 ± 0.007 p<0.05, Right PC left turn favorable outcome ΔF/F -0.02 ± 0.02 vs unfavorable outcome ΔF/F 0.04 ± 0.02 p<0.05, Left PC right turn favorable outcome ΔF/F -0.014 ± 0.008 vs unfavorable outcome ΔF/F 0.015 ± 0.008 p<0.05, Right PC right turn favorable outcome ΔF/F 0.002 ± 0.01 vs unfavorable outcome ΔF/F 0.02 ± 0.01 p=0.11 during replay, one-sided Wilcoxon signed-rank test with Benjamini-Hochberg correction for multiple comparisons.

Figure 5g: Mean Right PC ΔF/F - Mean Left PC ΔF/F for left turn Unfav - fav for closed loop -0.15 ± 0.06 vs replay 0.03 ± 0.04 p<0.05, Mean Right PC ΔF/F - Mean Left PC ΔF/F for right turn Unfav - fav for closed loop 0.1 ± 0.04 vs replay -0.01 ± 0.02 p<0.05, Mean Right PC ΔF/F - Mean Left PC ΔF/F for left turn Unfav - fav -0.15 ± 0.06 vs right turn Unfav - fav 0.1 ± 0.04 p<0.05 during closed-loop. Wilcoxon signed-rank test with Benjamini-Hochberg correction for multiple comparisons.

Figure 5h: Test accuracy score 5-fold cross-validation for decoding turn-outcome data 32.47 ± 1.6% vs shuffle 27.4 ± 1.1% p<0.01 for closed loop. Test accuracy score 5-fold cross-validation for decoding turn-outcome data 27.09 ± 2.3% vs shuffle 23.76 ± 0.4% p=0.08 for replay. Test accuracy score 5-fold cross-validation for decoding turn-outcome closed-loop data 32.47 ± 1.6% vs replay data 27.09 ± 2.3% p<0.05. Wilcoxon signed-rank test with Benjamini-Hochberg correction for multiple comparisons.

Figure S5e: Mean Right IO ΔF/F - Mean Left IO ΔF/F CW 0.62 ± 0.1 vs CCW -0.49 ± 0.09 p<0.01, Mean Right IO ΔF/F - Mean Left IO ΔF/F right optic flow 0.27 ± 0.05 vs left optic flow -0.22 ± 0.04 p<0.01, Wilcoxon signed-rank test with Benjamini-Hochberg correction for multiple comparisons.

Figure S5h: Test accuracy 5-fold cross-validation for decoding turn outcome data vs shuffle at 4 time points. -0.3 sec 0.31 ± 0.02 vs 0.29 ± 0.01 p=0.26, 0.3 sec 0.37 ± 0.02 vs 0.30 ± 0.01 p<0.001, 0.6 sec 0.47 ± 0.03 vs 0.29 ± 0.01 p<0.001, 1 sec 0.47 ± 0.04 vs 0.3 ± 0.01 p<0.001. Wilcoxon signed-rank test with Benjamini-Hochberg correction for multiple comparisons.

Figure S6d: Mean Right PC ΔF/F - Mean Left PC ΔF/F CW -1.02 ± 0.16 vs CCW 0.61 ± 0.09 p<0.001, Mean Right PC ΔF/F - Mean Left PC ΔF/F right optic flow -0.32 ± 0.05 vs left optic flow 0.12 ± 0.07 p<0.001, Wilcoxon signed-rank test with Benjamini-Hochberg correction for multiple comparisons.

Figure S6e: Test accuracy score 5-fold cross-validation for decoding turn outcome data vs shuffle at 4 time points. -0.3 sec 0.27 ± 0.01 vs 0.27 ± 0.01 p=0.57, 0.3 sec 0.29 ± 0.01 vs 0.28 ± 0.01 p=0.57, 0.6 sec 0.36 ± 0.02 vs 0.27 ± 0.01 p<0.01, 1 sec 0.40 ± 0.02 vs 0.27 ± 0.01 p<0.01. Wilcoxon signed-rank test with Benjamini-Hochberg correction for multiple comparisons.

Figure S6f: Mean turn bias after favorable outcome -0.008 ± 0.02 vs unfavorable outcome -0.069 ± 0.02 p<0.01, Mann-Whitney U test.

Figure 6e: Test accuracy score 5-fold cross-validation for decoding turn-outcome data 0.51 ± 0.04 vs shuffle 0.31 ± 0.02 p<0.05 for optic flow at 45°. Test accuracy score 5-fold cross-validation for decoding turn-outcome data 0.46 ± 0.04 vs shuffle 0.3 ± 0.01 p<0.05 for optic flow at 60°. Test accuracy score 5-fold cross-validation for decoding turn-outcome data (45°) 0.51 ± 0.04 vs data (60°) 0.46 ± 0.04 p=0.17 Wilcoxon signed-rank test with Benjamini-Hochberg correction for multiple comparisons.

Figure 6f: Test accuracy score 5-fold cross-validation for decoding turn-outcome data 0.37 ± 0.04 vs shuffle 0.29 ± 0.02 p<0.05 for optic flow at 45°. Test accuracy score 5-fold cross-validation for decoding turn-outcome data 0.39 ± 0.03 vs shuffle 0.3 ± 0.03 p<0.05 for optic flow at 60°. Test accuracy score 5-fold cross-validation for decoding turn-outcome data (45°) 0.37 ± 0.04 vs data (60°) 0.39 ± 0.03 p=0.81 Wilcoxon signed-rank test with Benjamini-Hochberg correction for multiple comparisons.

FigureS7e: Linear mixed-effects model, Mean (Right Pretectum Activity) - Mean (Left Pretectum Activity) *∼* Rank(Angle) + (1|fish): angle-rank slope = 0.040 ± 0.005, intercept = −0.088, p < 0.0001, n = 13 larvae.

## Acknowledgments

We thank all the members of the Portugues lab for constructive feedback and discussions throughout the project, especially Emanuele Paoli and Younes Farouj. We also thank Martin Haesemeyer and Michael Orger for comments on the manuscript. We would also like to thank the members of AS’s thesis advisory committee - Benjamin Judkewitz, Julijana Gjorgjieva, and Thomas Frank for their continued guidance throughout the project. This research was funded by the German Research Foundation (DFG) under Germany’s Excellence Strategy within the framework of the Munich Cluster for Systems Neurology (EXC 2145 SyNergy, identifier 390857198) and through Cornell University startup funds. AS was also partially funded by a scholarship from the Graduate School of Systemic Neurosciences, LMU.

## Author Contributions

AS and RP conceived and designed the study. AS performed the experiments, analyzed the data, and wrote the manuscript with inputs from RP.

## Data and code availability

All data described in this study, along with the analysis, simulation, and figure-generation notebooks, will be made publicly available upon publication. Whole-brain imaging data used in Figure 6 and Figure S7 was taken from Lavian, H. *et al.* Visual motion and landmark position align with heading direction in the zebrafish interpeduncular nucleus. *Nature Communications* **16.** doi:10.1038/s41467-025-66084-1 (2025)

## Supplementary Figures

**Figure S1:**
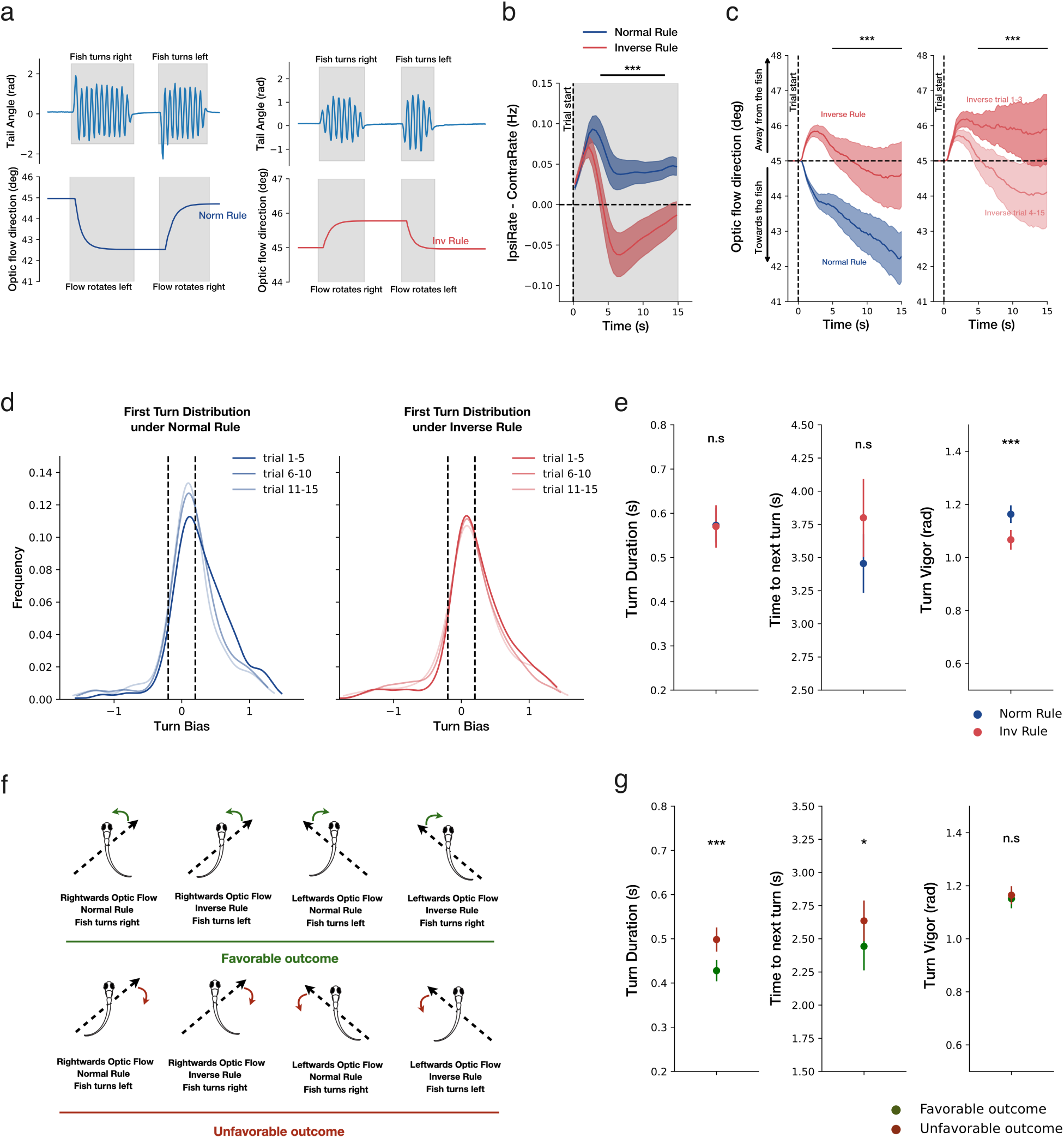
Behavioral dynamics during closed-loop normal and inverse rule trials. **a.** Example of tail trace (motor output) and associated change in the optic flow direction generated by the closed loop protocol during normal (left) and inverse rule (right). **b.** Difference between ipsilateral turn rate and contralateral turn rate across all trials during normal and inverse rule for the control group (top) and treatment group (bottom). **c.** Change in the optic flow direction during normal and inverse rule averaged across all trials (left) and change in optic flow direction during inverse rule averaged across trials 1-3 and trials 4-15 separately. **d.** Distribution of the first turn angle chosen under normal and inverse rule trials splits across first, middle and last 5 trials. **e.** Mean turn duration, time to next turn, and vigor for the first turn occurring in the trial during normal and inverse rule. **f.** All turns during normal and inverse rule trials are split into turns that generate a favorable outcome (optic flow rotates towards the fish) and unfavorable outcome (optic flow rotates away from the fish) (left) **g.** Mean turn duration, time to next turn, and vigor are shown for favorable and unfavorable outcomes. (right). All plots show mean and sem (shaded areas and error bars) across fish (n = 53 fish).

**Figure S2:**
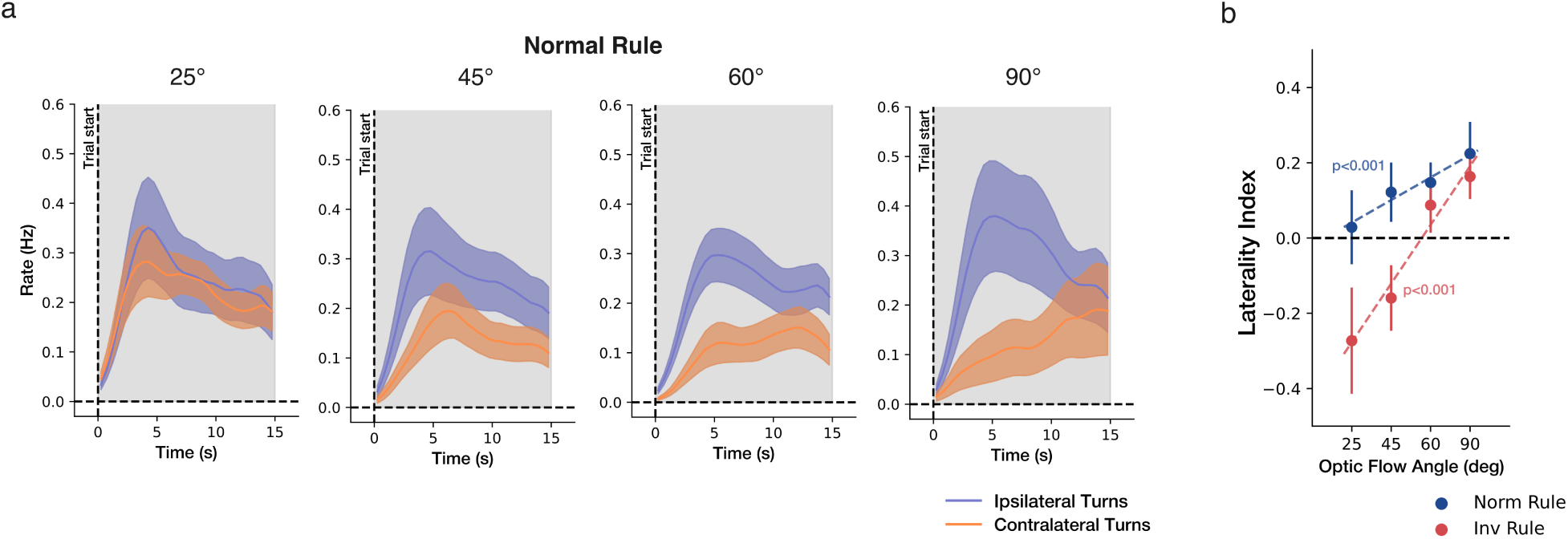
Normal feedback behavior for different optic flow angles. **a.** Mean ipsilateral(blue) and contralateral (orange) turn rate over the 15 s trial, averaged across trials, under normal rule, for each angle. **b.** Mean laterality index for each optic flow angle during normal and inverse rule trials. All plots show mean and sem (shaded areas and error bars) across fish (n = 18 fish).

**Figure S3:**
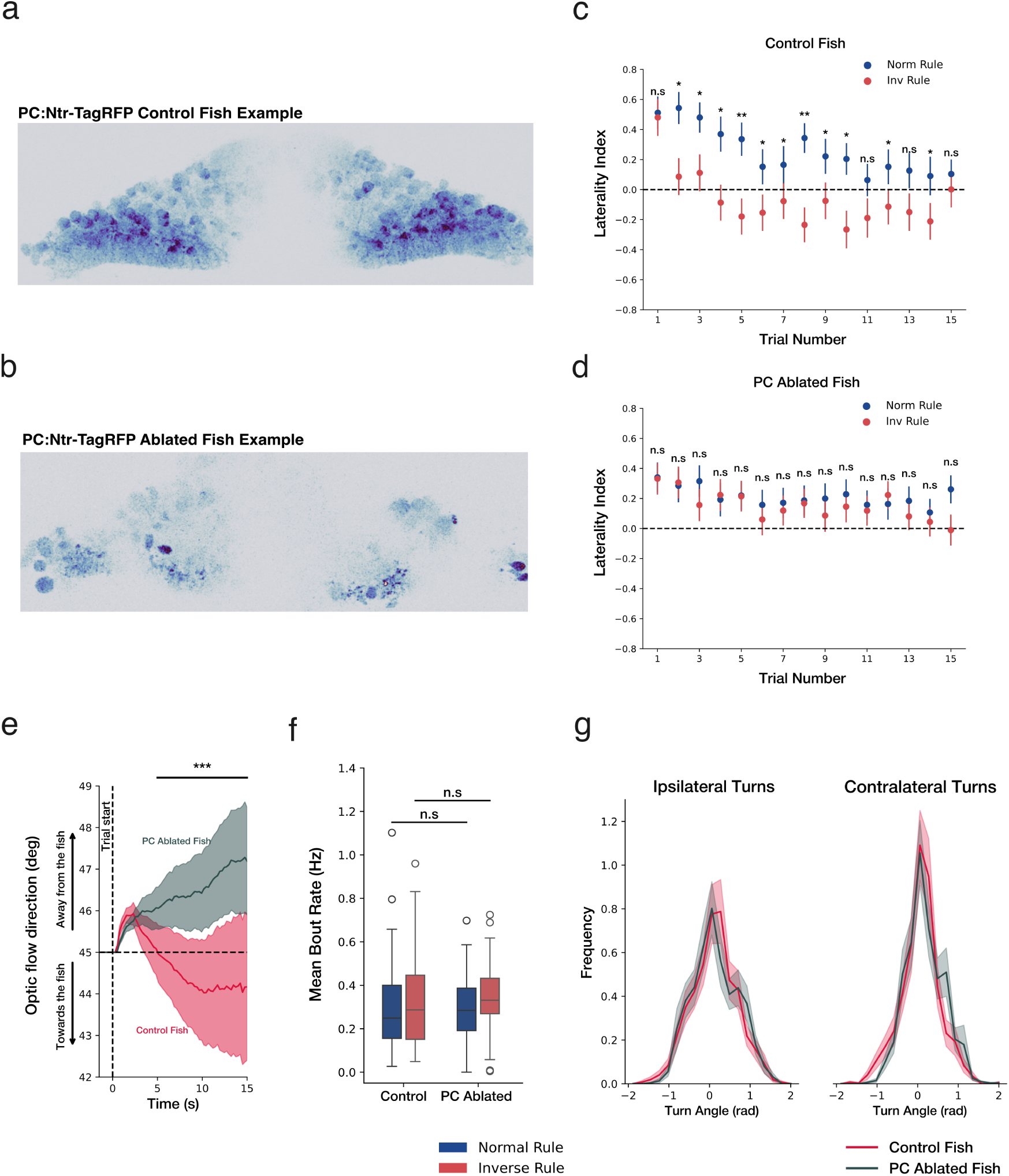
Effects of PC:Ntr-TagRFP ablation on closed-loop visuomotor behavior. **a.** Example confocal z-stack (max projection) of PC:Ntr-TagRFP positive larva before ablation. **b.** Example confocal z-stack (max projection) of PC:Ntr-TagRFP positive larva 24hrspostablation **c.** Trialwise laterality index for normal and inverse rule trials for control fish. **d.** Trialwise laterality index for normal and inverse rule trials for PC ablated fish. **e.** Change in the optic flow direction during inverse rule averaged across all trials for control and PC ablated fish. **f.** Mean Bout rate of control and PC ablated fish during norm and inv rule trials. **g.** Histogram showing turn angle distribution of the ipsilateral and contralateral turns for control and PC ablated fish. All plots show mean and sem (shaded areas and error bars) across fish (n = 21 control and 27 ablated fish).

**Figure S4:**
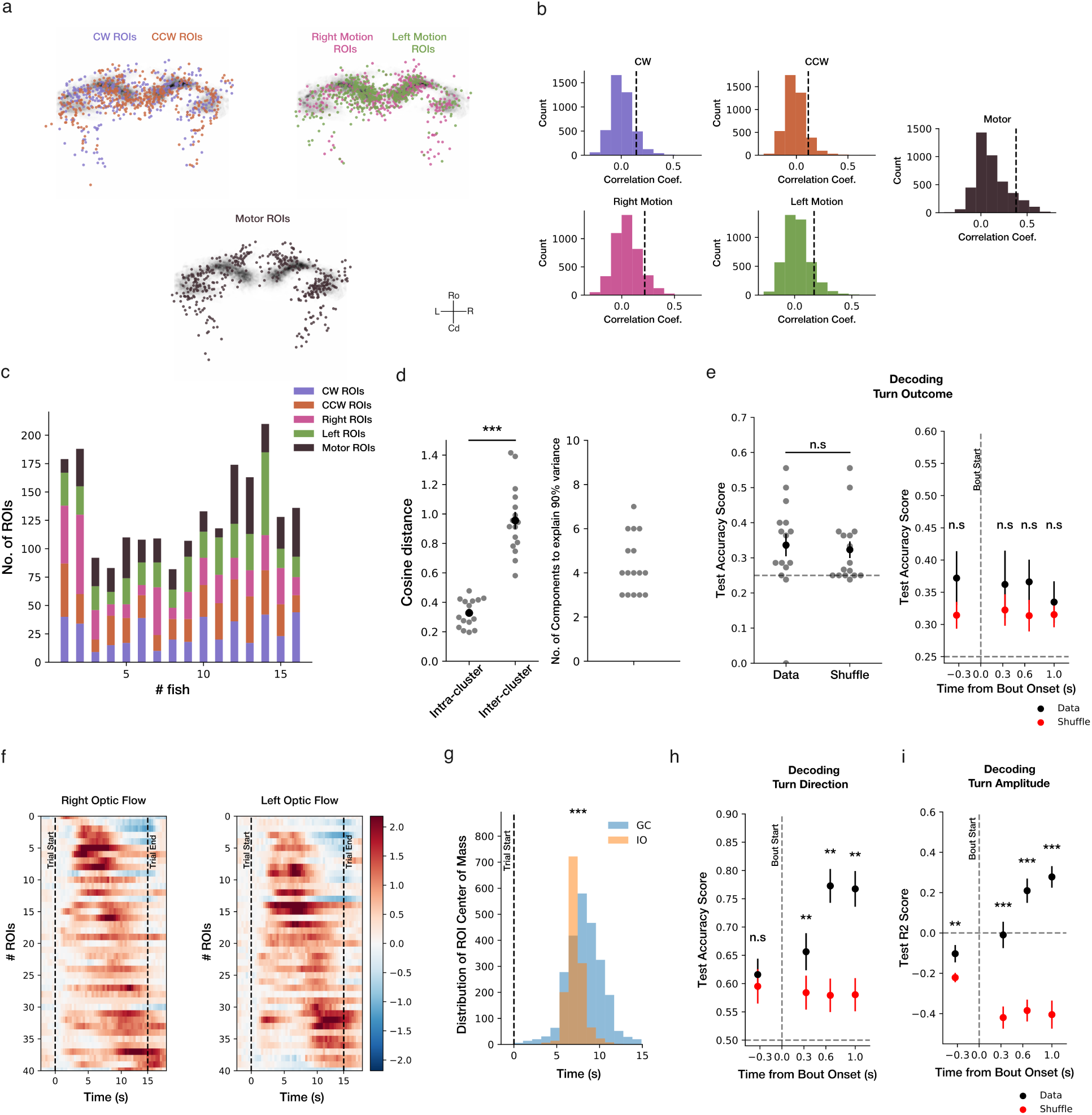
Functional organization and temporal dynamics of granule cell population activity. **a.** Anatomical maps of granule cell populations showing ROIs selectively correlated with clockwise (CW) and counterclockwise (CCW) rotation (top left), rightward and leftward optic flow motion (top right), and motor-related activity (bottom). **b.** Histograms showing the distribution of correlation coefficients between ROI activity and CW, CCW, right motion, left motion, and motor regressors across all fish. Dashed vertical lines indicate the top 10th percentile threshold used to assign ROIs to functional clusters. **c.**. Bar plot showing the number of ROIs assigned to each functional cluster across individual fish. **d.** Mean cosine distance between PCA activity trajectories computed within and between rightward and leftward optic flow-responsive clusters, quantifying the separability of sensory population dynamics. **e.** Decoding performance of a multiclass logistic regression model (5-fold cross-validation) trained to jointly predict turn outcome from granule cell population activity during closed-loop behavior in 1 sec period after bout onset(left) and at four distinct timepoints surrounding bout onset(right). **f.** Example fish showing temporally patterned sensory responses in granule cells during rightward optic flow (left) and leftward optic flow (right). ROIs are ordered according to response timing. **g.** Distribution of response half-maxima times for sensory-responsive ROIs in granule cell and inferior olive populations across all fish. **h.** Cross-validated test accuracy of a logistic regression classifier with 5-fold cross-validation trained to decode turn direction from motor-related granule cell activity at four distinct timepoints surrounding behavior. **i.** Cross-validated test R2 scores of a linear regression model with 5-fold cross-validation trained to decode turn amplitude from motor-related granule cell activity at four distinct timepoints. All plots show mean and sem (error bars) across fish (n=16 fish).

**Figure S5:**
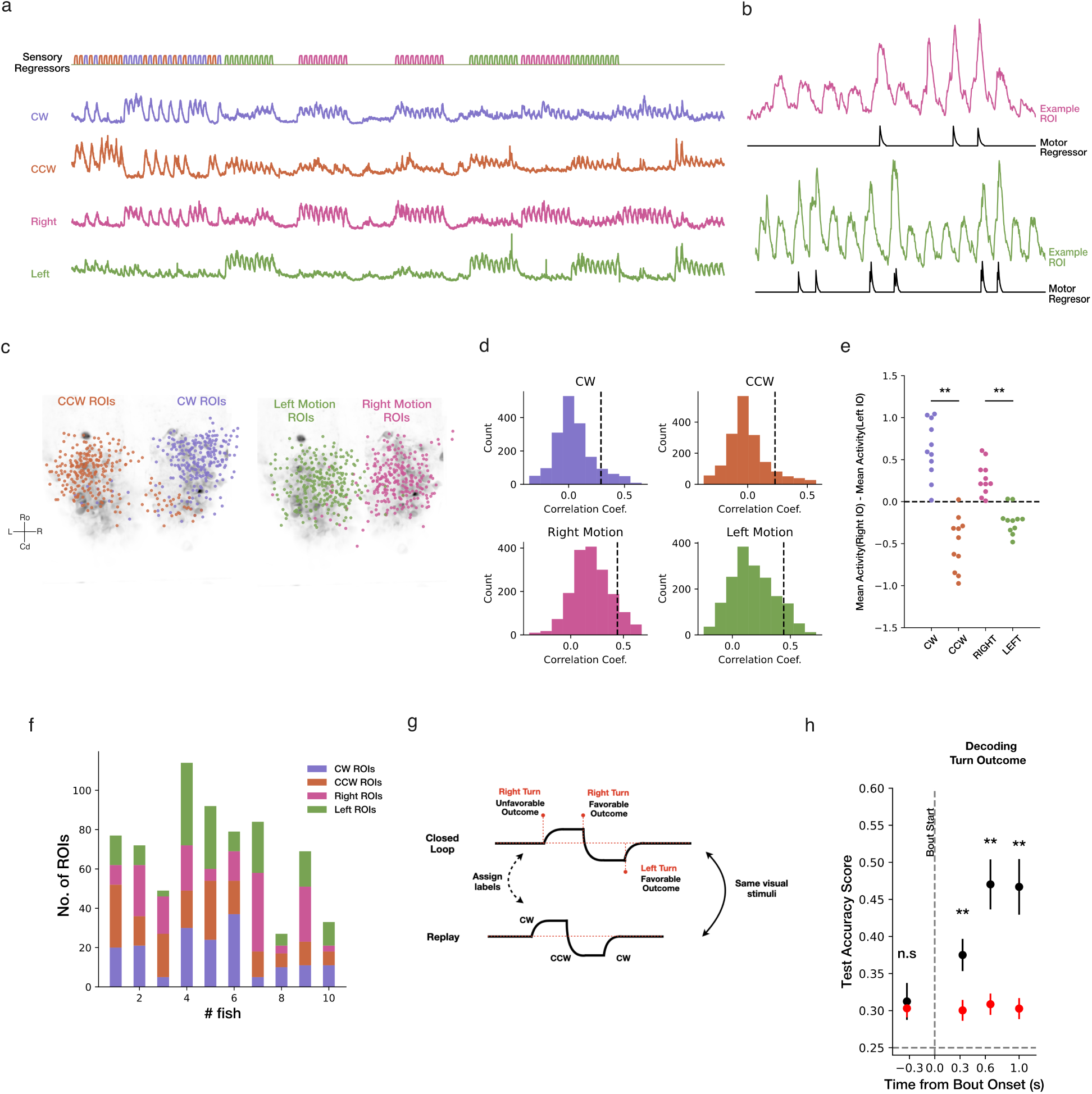
Characterization of IO population activity. **a.** Sensory clusters in IO activity for one example fish. **b.** Two example ROIs showing motor-induced activity in sensory IO clusters. **c.** Anatomical map of IO population showing ROIs selectively correlated with clockwise (CW) and counterclockwise (CCW) rotation (left), rightward and leftward optic flow motion (right). **d.** Histograms showing the distribution of correlation coefficients between ROI activity and CW, CCW, right motion, left motion regressors across all fish. Dashed vertical lines indicate the top 10th percentile threshold used to assign ROIs to functional clusters. **e.** Lateralization of IO sensory responses. **f.** Bar plot showing the number of ROIs assigned to each functional cluster across individual fish. **g**. Schematic illustrating assignment of turn direction and outcome labels during replay periods used for analysis of IO activity. **h**. Cross-validated test accuracy of a logistic regression classifier with 5-fold cross-validation trained to jointly decode turn outcome from ION activity at four distinct timepoints surrounding behavior. All plots show mean and sem (error bars) across fish (n=11 ION fish).

**Figure S6:**
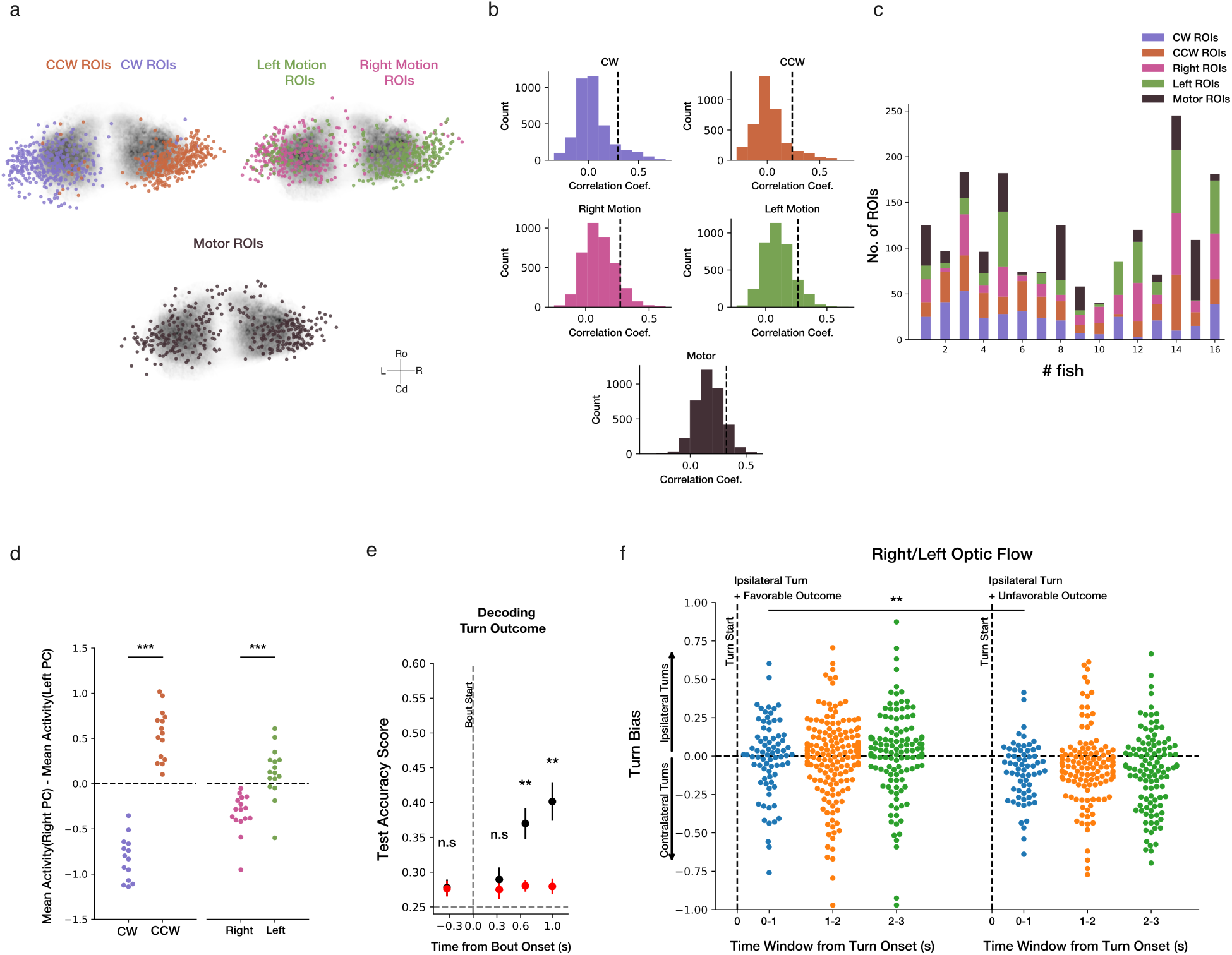
Characterization of PC population activity. **a.** Anatomical maps of Purkinje cells showing ROIs selectively correlated with clockwise (CW) and counterclockwise (CCW) optic flow (top left), rightward and leftward optic flow motion (top right), and motor-related activity (bottom). **b.** Histograms showing the distribution of correlation coefficients between ROI activity and CW, CCW, right motion, left motion, and motor regressors across all fish. Dashed vertical lines indicate the top 10th percentile threshold used to assign ROIs to functional clusters. **c.** Bar plot showing the number of ROIs assigned to each functional cluster across individual fish. **d.** Lateralization of PC sensory responses. **e.** Cross-validated test accuracy of a logistic regression classifier with 5-fold cross-validation trained to jointly decode turn outcome from PC activity at four distinct timepoints surrounding behavior. **f.** Turn bias in successive 1-s windows following a turn that received a favorable or unfavorable outcome, up to 3 s after the bout. All plots show mean and sem (shaded areas and error bars) across fish (n=16 PC fish).

**Figure S7:**
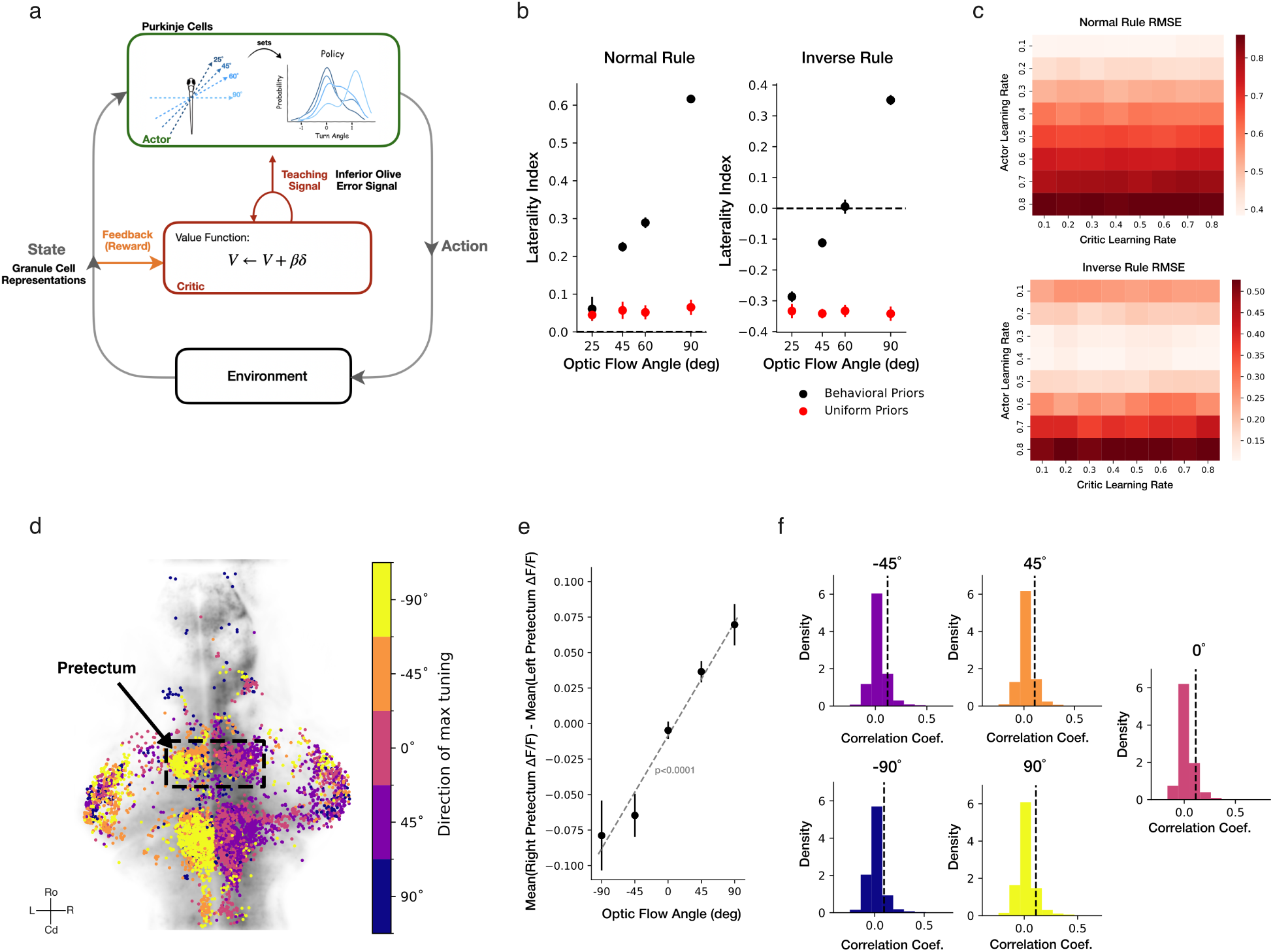
Model controls and pretectal representation of optic flow. **a.** Schematic of the actor–critic model. **b.** Laterality index of the agent across trials under normal and inverse rule, for a policy initialized with the measured innate (behavioral) prior versus a uniform prior. **c.** Root-mean-square error (RMSE) between experimentally observed and simulated behavioral correction across optic-flow angles, evaluated over a grid of learning-rate parameters. **d.** Whole-brain map from a representative larva showing ROIs in the top-10 percentile of correlation with one direction of optic flow. **e.** Difference between mean right and mean left pretectal activity across optic flow angles with linear mixed-effects model fit **f.** Distribution of correlation coefficients between pretectal ROIs (pooled across larvae) and each optic flow direction. Dotted lines indicate the top-10 percentile threshold. All plots show mean and sem (error bars) across fish (n=18 fish for normal rule turn distribution, 7 fish for ION activity at 45° and 60°, 7 fish for PC activity at 45° and 60°, and 13 fish for pretectum activity). The pretectum data in d, e and f are reanalyzed from [62].

